# Illusory tactile movement crosses arms and legs and is coded in external space

**DOI:** 10.1101/2021.05.20.445020

**Authors:** Marie Martel, Xaver Fuchs, Jörg Trojan, Valerie Gockel, Boukje Habets, Tobias Heed

## Abstract

Humans often misjudge where on the body a touch occurred. Theoretical accounts have ascribed such misperceptions to local interactions in peripheral and primary somatosensory neurons, positing that spatial-perceptual mechanisms adhere to limb boundaries and skin layout. Yet, perception often reflects integration of sensory signals with prior experience. On their trajectories, objects often touch multiple limbs; therefore, body-environment interactions should manifest in perceptual mechanisms that reflect external space.

Here, we demonstrate that humans perceived the cutaneous rabbit illusion – the percept of multiple identical stimuli as hopping across the skin – along the Euclidian trajectory between stimuli on two body parts and regularly mislocalized stimuli from one limb to the other. A Bayesian model based on Euclidian, as opposed to anatomical, distance faithfully reproduced key aspects of participants’ localization behavior.

Our results suggest that prior experience of touch in space critically shapes tactile spatial perception and illusions beyond anatomical organization.

## Introduction

Our everyday life is defined by interactions between body and environment. Action and perception work together to create a loop: we perceive to act and act to perceive (Ernst & Bülthoff, 2004; Friston, 2010; Gibson, 1966, 1979; Neumann & Prinz, 1990; Witt & Riley, 2014; Wolpert & Ghahramani, 2000). Through this loop, our perception of the world and of our own body in space is shaped by experience. The brain interprets incoming sensory input and employs assumptions, or expectations, to compensate for uncertainty that is due to sensory noise. The resulting percept is often a compromise between the actual sensory input and experience-based expectations (Goldreich, 2007; Goldreich & Tong, 2013; Körding & Wolpert, 2004; Shams & Beierholm, 2010). The compromise between sensory input and expectation can evoke perceptual illusions, which usually express the brain’s expectations overriding sensory mismatch (e.g. Brown et al., 2013; Hayward, 2015). For instance, touch is not always perceived at its true location but is ascribed elsewhere to better “fit” with a perceptual expectation.

The famous cutaneous rabbit effect is a prime example of a perceptual illusion: A rapid sequence of tactile stimuli, presented at two positions on the skin several centimeters apart, evokes the percept of a stimulus sequence that gradually moves along the stretch of skin between the truly stimulated positions (e.g. Geldard & Sherrick, 1972; Trojan et al., 2010). A common explanation for this percept is that it arises from the brain’s assumption that stimuli that occur close to each other in time and space originate from one common source (Goldreich, 2007; Goldreich & Tong, 2013; Tong et al., 2016); in the case of the cutaneous rabbit, the common source may be a hypothetical rabbit that hops across the skin, giving the illusion its name. This idea is elegantly captured by a Bayesian observer model that posits that the brain uses a low velocity prior, that is, a generalized expectation that successive stimuli are caused by a common object that moves continuously along the skin at a speed typically encountered in real life. The model gives higher relative weight to the prior the higher spatial uncertainty about the sensory input is, which can ultimately result in a percept that reflects previous experience more than the present input.

How could a prior favoring the rabbit-like percept arise? On many occasions, we perceive things moving along the skin, creating an orderly sequence of tactile input. Every occurrence of a moving object along the skin updates and strengthens the brain’s expectation that this type of stimulation pattern is caused by one common, persistent object. Some studies, especially in the visual modality, have shown that exposure to altered rules, such as stimulus probabilities and the contingency between different sensory inputs, change our perceptual priors (Adams et al., 2004; Barnett-Cowan et al., 2018; Di Luca & Rhodes, 2016). For instance, visual patches are typically perceived as convex when they are brighter on the top half. Yet, this judgment can change with visuo-haptic training, with participants effectively modifying their “light-from-above” prior (Adams et al., 2004). Generally, then, experience-shaped priors reflect stimulation patterns that arise from regularities in the world, such as how our body in space normally encounters moving objects.

Critically, the use of external space within the framework of moving objects is not straightforward in the tactile system. Our tactile sensors lie in the 2D sheet that is our skin, and this anatomically based 2D reference frame must be converted into an external reference frame that reflects where the touch happened in space. This transformation entails combining skin location with posture, and, when an object moves across several body parts, taking into account the spatial relationships between them (Heed et al., 2012; Shore et al., 2002; Yamamoto & Kitazawa, 2001). Hence, anatomical **and** external space are firmly linked in the tactile system (for review see Badde & Heed, 2016; Longo et al., 2010; Medina & Coslett, 2010; Tamè et al., 2019).

Given the central role of spatial priors in the Bayesian observer model and given that these priors reflect previous experience gained through environmental interactions – that is, interactions that take place in external space –, it may strike as surprising that, until now, the model has always been applied to anatomical space. Similarly, explanations of the cutaneous rabbit effect have almost exclusively focused on the illusion as a phenomenon related to anatomical, and not to external spatial processing. For example, early studies proposed low-level interactions in the skin and neurons in primary somatosensory regions (S1) as the effect’s origin (Geldard, 1975, 1985; Geldard & Sherrick, 1972, 1986). Several findings corroborated this view. For instance, the illusory rabbit was reported to not cross the body midline (Geldard, 1982, 1985; but see Eimer et al., 2005; Trojan et al., 2010 for opposing results); the size of the illusion appeared to covary with the receptive field size of the presumed corresponding neurons in S1 (Geldard & Sherrick, 1983); and the only neuroimaging study that has investigated the rabbit phenomenon (Blankenburg et al., 2006) observed similar activity in S1 when a location was truly stimulated and when it was perceived due to the rabbit illusion. In fact, the illusory percept of a location in the middle between the two truly stimulated locations also lay at an intermediate S1 position, suggesting a parallel of the perceived stimulus pattern and neural S1 activity.

If the suggestion were true that the cutaneous rabbit effect is bound to anatomy, the rabbit illusion would be limited to 2D skin space and, in extension, to neighboring neural regions in S1. Accordingly, it could not occur across distant body parts, such as an arm and a leg. Yet, the idea that tactile spatiotemporal illusions are bound to the body, or anatomical space, is at odds with the proposition that they are based on expectations about interactions with the environment: Objects in the environment move in external space. Only under very specific circumstances are anatomical and external space aligned. For example, when sitting on the grass at a picnic, ants will crawl from one finger to the next, and from the hand onto the leg when you’re resting your arms on your lap. They will not only walk along single limbs. Hence, they create sensory signals in receptive fields that are remote in anatomical space. Establishing a prior based on neighboring receptive fields (i.e., anatomical space) would not faithfully reflect these experiences. Thus, it is insufficient to consider only anatomical space to tune the brain to movement in space. Accordingly, such a strategy would not be an optimal adaptation to interacting with the environment.

Some studies have indirectly related the rabbit illusion to external space. For instance, a rabbit percept was found to cross the midline at the abdomen, implying involvement of both hemispheres (Trojan et al., 2010). Stimuli were also perceived as if moving along a stick held between the index fingers while only the fingers had been stimulated, showing that the cutaneous rabbit effect could be experienced outside the body (Miyazaki et al., 2010). Last, it has been observed that stimuli presented on both forearms, that is, limbs represented in S1 of different hemispheres, attract each other (Eimer et al., 2005). However, all these studies applied the rabbit paradigm to homologous body parts. Accordingly, across-limb rabbit percepts may be related to interhemispheric connectivity between homologous body regions of the two body halves, effectively corroborating the dependence of the rabbit percept on anatomically guided neural organization. Finally, two studies have reported rabbit effects between different fingers of one hand (Trojan et al., 2014; Warren et al., 2010). Yet, the fingers, again, activate neurons in neighboring areas of S1, and so rabbit jumps between fingers may not be considered strong evidence for illusory movement in external space. Overall, then, a growing number of studies have casted doubt on the original proposal that illusory localization, as observed in the rabbit illusion, is solely determined by anatomical criteria, but none of them has provided strong and explicit evidence for a determining role of external space in the priors reflected by the illusion. Here, we present direct evidence for an important role of external spatial coding in the rabbit percept. We presented stimulus sequences known to elicit the rabbit illusion to single limbs as well as across two limbs placed next to each other. Stimuli presented to the different limbs were close together in space, because participants placed their limbs next to each other. Under the assumption that tactile stimuli are evaluated in 3D space, our setup made it plausible that the sequences’ tactile events were caused by a common origin. The rabbit effect was evident across the two arms, and across an arm and a leg of the same and of opposite body sides. Participants indeed reported that stimuli were distributed along the hypothetical trajectory between the true stimulus locations, whether the sequence was on one limb or distributed across two limbs. Moreover, participants sometimes reported displaced stimuli on the incorrect limb, implying that perceptually displacing stimuli in space also affected how these stimuli were assigned to a body part. Together, these findings suggest that the cutaneous rabbit effect involves an external-spatial process that can evoke not only spatial distortions but, in addition, categorical errors about the limb on which a stimulus seems to have occurred.

## Material and methods

No part of the study procedures/analyses was pre-registered prior to the research being conducted. We report how we determined our sample size, all data exclusions, all inclusion/exclusion criteria, whether inclusion/exclusion criteria were established prior to data analysis, all manipulations, and all measures in the study

### Experiment 1

#### Participants

We aimed to acquire 20 participants based on previous studies using similar methods that have recruited 15 to 20 (Eimer et al., 2005; Trojan et al., 2010, 2014). Of 21 acquired data sets, we had to exclude two, which left us well in the range of previous studies (n = 19, 13 females, mean age = 25 years, SD = 6). One participant was excluded because their data pattern suggested that two stimulators may have been switched during set-up, and one was excluded because (s)he was unable to maintain the required postures. We determined handedness by splitting Edinburgh Handedness Inventory scores (EHI; Oldfield, 1971) into [−100; −61] for left-handedness, [−60; 60] for ambidextrousness, and [61; 100] for right-handedness. One participant was left-handed, three were ambidextrous, and the remaining 15 participants right-handed. All participants reported normal or corrected-to-normal vision and were free of any known neurological, visual, motor or tactile disorders. Participants were naïve about the purpose of the experiment and gave written informed consent prior to the experiment. They either participated without compensation or received course credit. The local ethics committee of Bielefeld University approved the experiment.

#### General procedures

Participants sat at a desk, in front of a 23.8-inch (52.7 x 29.6 cm) computer monitor (Dell SE2417H, Dell, Round Rock, TX, USA) at an approximate distance of 50-75 cm. The picture of a limb was displayed on the screen in the same posture as the participant’s limb(s) (Figure 1; Flach & Haggard, 2006). The image files were 1920 x 1028 pixels, corresponding to the screen’s resolution. When a single arm was displayed, it was depicted from the elbow to the tip of the third digit and covered ∼1700 px in width. This corresponded to a life-like size of ∼46 cm. When two limbs were shown, each arm had to be depicted slightly smaller so that both arms would fit the screen; each arm then covered ∼1350 px or ∼37 cm. The image was filled with white color around the arm(s). All images are shown in Table 1. We used mirror images for limb configurations with the left and the right body side (see Table 1). Participants wore a cape both during setup and testing that prevented view of their stimulated limb(s) to reduce possible visual anchors (Trojan et al., 2010). Before each trial, there was a random inter trial interval, ranging between 1000 and 1500ms, during which a message on the screen indicated “Please focus on the touch”. If participants indicated that they did not feel one of the stimuli, the experimenter pressed a key to repeat the trial at the end of the block. This happened rarely (11 of 65082 trials = 0.02%).

**Figure 1.**
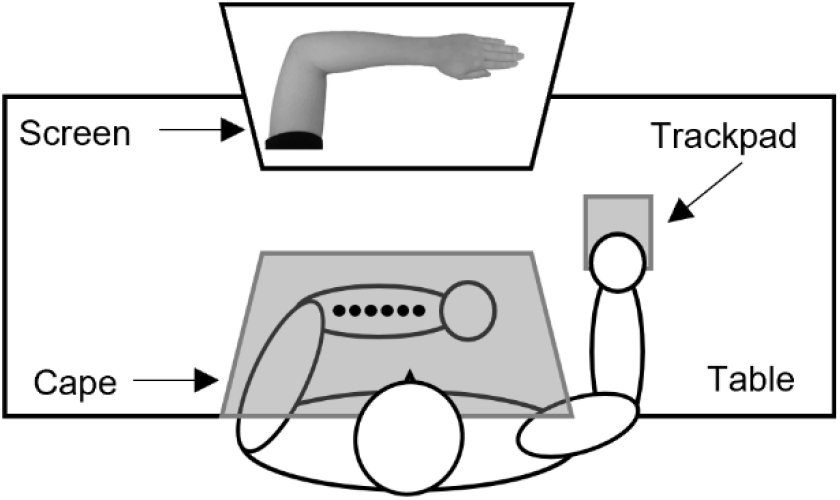
Experimental setup. The schematic depicts the configuration “Left Single Arm” (see the following section for details of the other configurations).

**Table 1.**
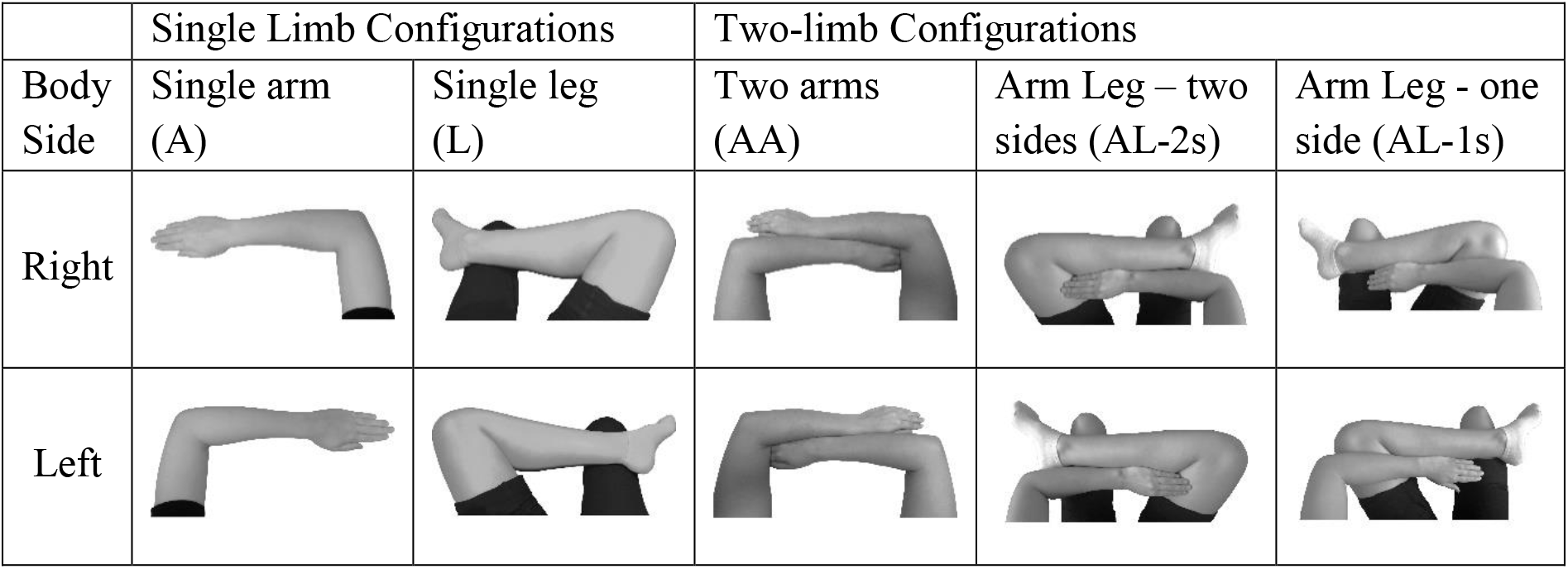
Images presented on the screen depending on limb configurations. Participants performed each of the five configurations in a pseudo-randomized order (participants always performed all the single or the two-limb configurations first). Half of the participants performed with their right body side, half with the left.

Stimulation involved either one or three stimuli presented to participants’ limb(s). Participants then reported where they had felt each stimulus by clicking on the corresponding location on the image on the screen using a trackpad. This method was similar to the one used by Flach & Haggard (2006) and allowed a direct measure of localization errors. The trackpad was always placed so that it could be comfortably operated with a free hand, and whenever possible, with the dominant hand. The experiment with its five limb configurations lasted ∼3 hours. As seen in Table 1, these configurations were divided in single limb configurations, involving either an arm or leg alone, and two-limb configurations. Two-limb configurations comprised an arm and a leg of the same or different body sides, or two arms. Half of the participants started with single limb configurations, the other half with two-limb configurations (order of the configurations was then randomized). There was a short break of ∼10 min between each limb configuration. For each configuration, participants first performed a block in which they localized individually presented stimuli (“single-location task”). Then, different types of three-stimulus sequences (described below) were presented in randomized order (“three-location task”). Participants localized the three consecutive tactile stimuli in perceived order. In half of our sample, we stimulated the left body side in single-limb configurations, and the right side in the other half. At the end of the session, we asked participants to judge stimulus intensity on the arm or leg on a visual scale from zero (= no sensation) to ten (= very strong sensation). One stimulator was attached to the middle of each limb and participants rated its intensity. Participants judged the stimuli on the arm significantly stronger than on the leg (5.2 ± 2.0 vs. 3.0 ± 1.3; SE = .448, t = −4.93, p = .039).

#### Tactile stimuli

Eight solenoid vibrotactile stimulators (Dancer Design, St. Helens, UK; diameter 1.8 cm) were wired to a custom-built control unit (Neurocore, Hamburg, Germany) that was connected to the computer. The Neurocore triggered the stimulators with sub-millisecond precision. The experiment was controlled by the software PsychoPy (v1.85.6; Peirce, 2007) and triggered stimulus sequences on the Neurocore. Thus, timing inaccuracies induced by the Operating System may have affected the precision of the sequence start – which is of no interest –, but did not affect the timing of the three stimuli relative to each other, which is critical for the rabbit illusion setup. Tactile stimuli consisted of 200 Hz sine waves presented for 45 ms by driving stimulators with a fixed output power for all stimuli. Participants wore headphones that played white noise to cancel any auditory noise from the stimulators. We determined stimulator positions individually. We measured participants’ arm and leg length and marked the midpoint between wrist and elbow, as well as between ankle and knee. We used these lines to position casings that contained the stimulators and served as stencils for their placement. Stimulators were spaced 1.5 cm from border to border (or 3.3 cm from center to center, given their 1.8 cm diameter). This setup resulted in a center-to-center distance of ∼10 cm for any chain of 4 stimulators. Once positioned, the stimulators were taped to the skin with adhesive tape rings and the casing removed. Six stimulators were used for the single limb configurations (see Figure 2A), and eight for the two-limb configurations (see Figure 2B). For analysis and reporting, we numbered stimulators from 1 to 8 in ascending order from left to right (see example in Figure 2B). This numbering scheme was defined externally, not anatomically. Therefore, when we specify a particular stimulus pattern, it defines an identical sequence in space in all two-limb configurations. For example, in configuration AL-1s (left arm and left leg), the proximal-distal direction is from left to right for both limbs, but for configuration AL-2s it is proximal-distal for the arm and distal-proximal for the leg (see Figure 2B), but this difference is irrelevant for the way we speak about sequences.

**Figure 2.**
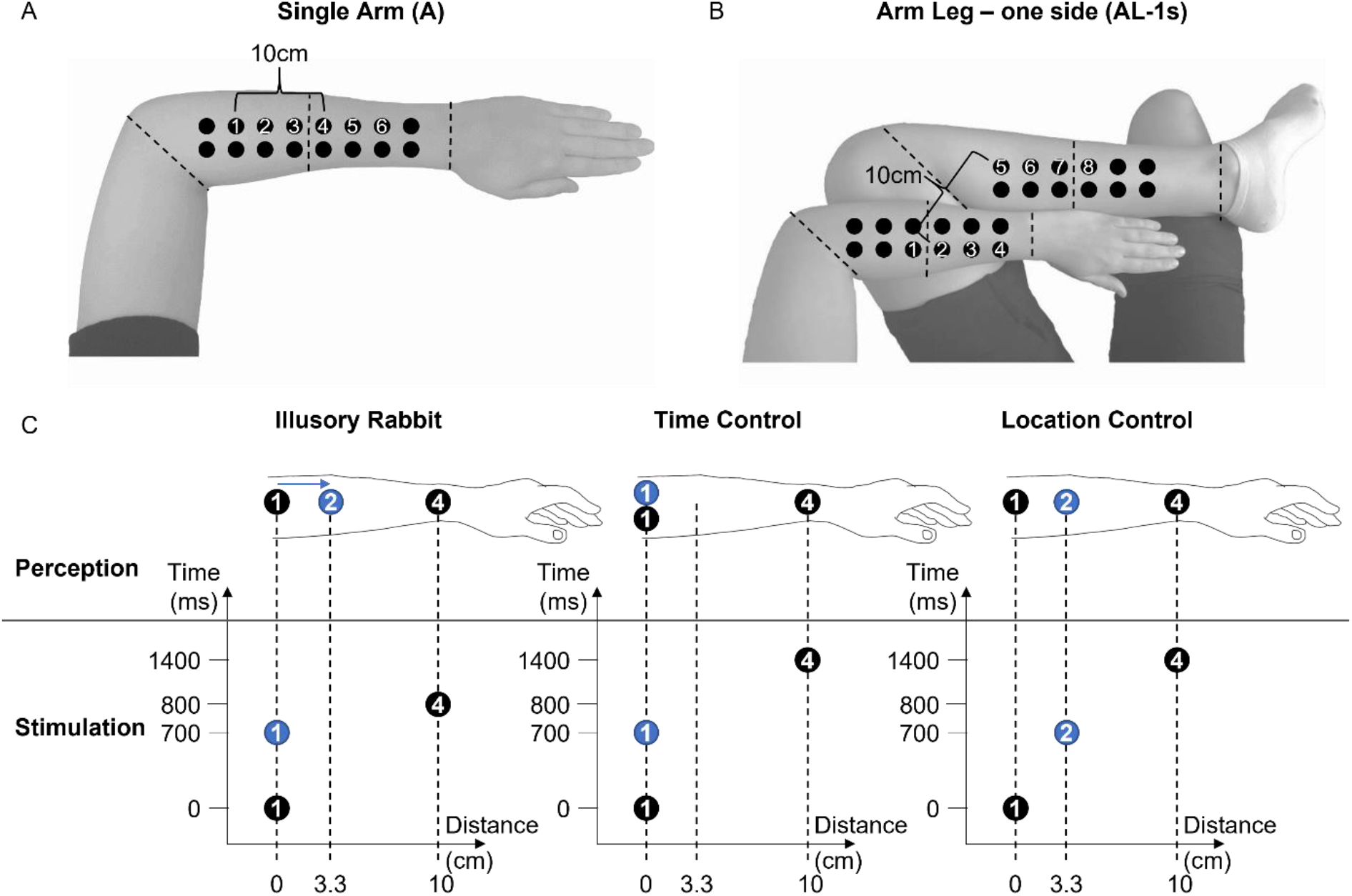
Stimulator positions for A/the single arm (A) and B/the arm leg – one side (AL-1s) configuration when stimulating the left body side. The positions were similar in the other limb configurations. We used only the 8 numbered stimulators; all others were attached only to obscure the relevant locations to participants at all times. In all configurations, the distance between the first and last stimuli was always 10 cm (e.g. stimulus positions 1-1-4 or 7-7-4). The dashed lines represent the landmarks used to place the stimulators (distance elbow/wrist or knee/ankle and their midpoint, see Methods). **(C) Schematics of the three-stimulus sequences** (Goldreich, 2007). Each panel schematically depicts one sequence. Numbered circles represent the three stimuli, with the numbers indicating the temporal order of the sequence (and matching the true number of the stimulator in panels A and B). In illusory rabbit and time control trials, the first and second stimuli occurred on the same skin location (same vertical dashed line). In the location control trials, the second stimulus was in a different location between the first and last stimulus (3.3 cm from the first stimulus in the example of the depicted 1-2-4 sequence); in two-limb configurations, the second stimulus could be on the same limb (e.g. 2-3-5 in panel B) or not (e.g. 4-6-7 in panel B). The first and third stimuli were always 10 cm apart. The Stimulus Onset Asynchrony (SOA) between the first and second stimuli was always 700 ms. The SOA between the second and third stimuli was either 100 ms or 700 ms, depending on stimulus sequence, as indicated on the y-axis. The resulting perception is indicated on the schematic drawing of the arm above the axis grids: in the illusory rabbit, the perception of the second stimulus is often shifted towards the third (as indicated by the blue arrow). In the two other sequences, the second stimulus is usually perceived near its true location.

Around the stimulators relevant for our sequences, we attached additional, inactive stimulators as dummies to obscure which stimulators would be active during the experiment. There were ten dummy stimulators for the single limb postures and twelve for the two-limb (Figure 2A,B). In two-limb configurations, we chose a positioning of limbs and stimulators that avoided spatially incoherent sequences. When the stimulation sequence moved from left to right, the last stimulus was presented on the right relative to the first one, and vice versa for right to left sequences. For this reason, stimuli 4 and 5 in Figure 2B were always vertically aligned, with stimulus 1-4 on the left side and stimulus 5-8 on the right side. Exploration of possible postures during the planning of our experiments had indicated that other combinations of postures and stimulus locations could confuse participants.

#### Stimulus sequences

In the first task, we used only single locations (“single-location task”). In the second task, we used three different stimulus sequences: illusory rabbit sequences, time control sequences, and location control sequences (“three-location task”). All trials of the three sequence types were presented in one block in randomized order. In all sequences, the Euclidian distance between the first and third stimuli was 10 cm, and the three sequence types differed only with respect to the second stimulus. Illusory rabbit sequences were smaller in number as compared to the time control and location control sequences to obscure the aim of the experiment.

The *single location* consisted of one stimulus per trial, at any of the six or eight possible locations in a random order (respectively for numbers 1-6 for the single or 1-8 for the two-limb configurations, see Figure 2A,B). Each position occurred six times so that this block consisted of 36 trials for each single limb configuration and 48 trials for each two-limb configuration. It served as a brief sanity check for participants’ ability to differentiate and localize the different stimulus positions when no other stimuli are present that might trigger spatiotemporal integration, and to familiarize participants with the localization procedure in the absence of any perceptual illusion effects.

The *illusory rabbit* sequences consisted of three stimuli per trial, the first two at one location and the last at another 10 cm away (Trojan et al., 2010). Stimulus Onset Asynchrony (SOA) was 700 ms between first and second stimulus, and 100 ms between the second and the third (Eimer et al., 2005; Figure 2C). These stimulus patterns are known to elicit the rabbit illusion, which is defined as perceiving the second stimulus as displaced towards the location of the third stimulus. We used six patterns for the single limb configurations (1-1-4, 2-2-5, 3-3-6, 6-6-3, 5-5-2, 4-4-1, see Figure 2A), and ten for the two-limb configurations (right body side: 1-1-4, 4-4-1, 8-8-5, 5-5-8, 1-1-6, 6-6-1, 2-2-7, 7-7-2, 3-3-8, 8-8-3; left body side: 1-1-4, 4-4-1, 8-8-5, 5-5-8, 2-2-5, 5-5-2, 3-3-6, 6-6-3, 4-4-7, 7-7-4; see Figure 2B). Left-to-right and right-to-left sequences were balanced. In one-limb configurations (A, L), all three stimuli occurred on the respective limb. In two-limb configurations, we presented cross-limb trials, but interspersed them with single-limb sequences with 3 stimuli on the same limb (first four patterns above). Mixing single-and cross-limb sequences served to prevent participants from habitually reporting stimuli on the limb stimulated second. Moreover, they allowed us to ascertain that the illusion emerges with its typical, one-limb stimulus layout even if other limbs are task-relevant, and served as sanity check that the illusion occurs normally also in our extended, rather complex setup. We report these single limb results in Supplementary Information (Table S1) but did not include them in the main analyses. Each sequence pattern occurred five times, so that this condition comprised 30 trials for single limb configurations and 50 trials for two-limb configurations (20 single limb and 30 cross-limb trials); sequences were fully randomized.

The *time control* sequences, too, consisted of three stimuli per trial. They were designed as a control condition for the illusory rabbit sequences and stimulated the same positions, but with a 700 ms SOA between all stimuli (Figure 2C). These sequences served as controls, as previous work has established that displacement of the second stimulus, indicative of a rabbit illusion is strongly reduced (even if not completely obliterated) with this stimulus timing, compared to the fast, 100 ms SOA of the illusory rabbit trials (e.g. Trojan et al., 2010). Stimulus patterns, stimulator positions, and sequence repetitions were as for the illusory rabbit sequences. The *location control* sequences were designed as another control sequence and again consisted of three stimuli per trial with 700 ms SOAs between all stimuli. However, these sequences stimulated three distinct locations, with the second stimulus located between the first and third (Figure 2C). We included these sequences so that participants would occasionally perceive three locations in other sequences than the illusory rabbit. This further obscured the aim of the experiment and avoided that participants were biased against reporting the second stimulus at intermediate locations. Single limb configurations comprised four patterns (1-2-4, 6-5-3, 4-2-1, 3-5-6; see Figure 2B), as did two-limb configurations (right body side: 1-7-6, 3-2-8, 6-7-1, 8-2-3; left body side: 2-3-5, 4-6-7, 5-3-2, 7-6-4; see Figure 2B). Each pattern occurred five times, so this block consisted of 20 trials for each limb configuration.

#### Data processing

We pre-processed and analyzed data with R (v3.6.1; RStudio v1.1.442; R Core Team, 2018). The coordinates of each localization response given by participants were recoded so that data were coded as if all participants had performed all configurations with their left body side, and the AL-2s configuration with their left arm and right leg; recoding allowed pooling all participants into one analysis while discounting stimulation side. We computed the Euclidean distance between the reported locations of the first and the third stimulus and projected the second perceived stimulus on the straight line between the first and third (Trojan et al., 2010, 2014, 2019). We computed the percentage of shift of the second stimulus toward the third (see Figure 3). Participants sometimes confused the order or direction of the stimuli, for instance by reporting positions of stimuli 1, 3, 2 instead of 1, 2, 3. This kind of error has been frequently observed by others at similar rates (Asai & Kanayama, 2012; Trojan et al., 2014). We excluded trials with order or direction errors (23% of all trials) with a tolerance distance margin of 10%, resulting in the remaining trials having a proportion of displacement between −10% and 110%. We considered the rabbit illusion to have been present if the displacement between the first and second stimulus was more than 10%. The 10% threshold is arbitrary, but similar conclusions resulted when we adopted other thresholds such as 5%; see Table S2 & Figure S1). With the 10% threshold, all displacements in the range of −10% to 10% were treated as if participants had reported the first and second stimuli in the same location and, thus, that the rabbit illusion had not occurred in the respective trial. Similarly, displacement of 100% would indicate that the second and third stimuli were perceived at the same location. For the two-limb configurations, we used image masks to determine whether the second stimulus had been reported on the limb on which it had been presented, or whether it had been assigned to the other limb (Figure 3).

**Figure 3.**
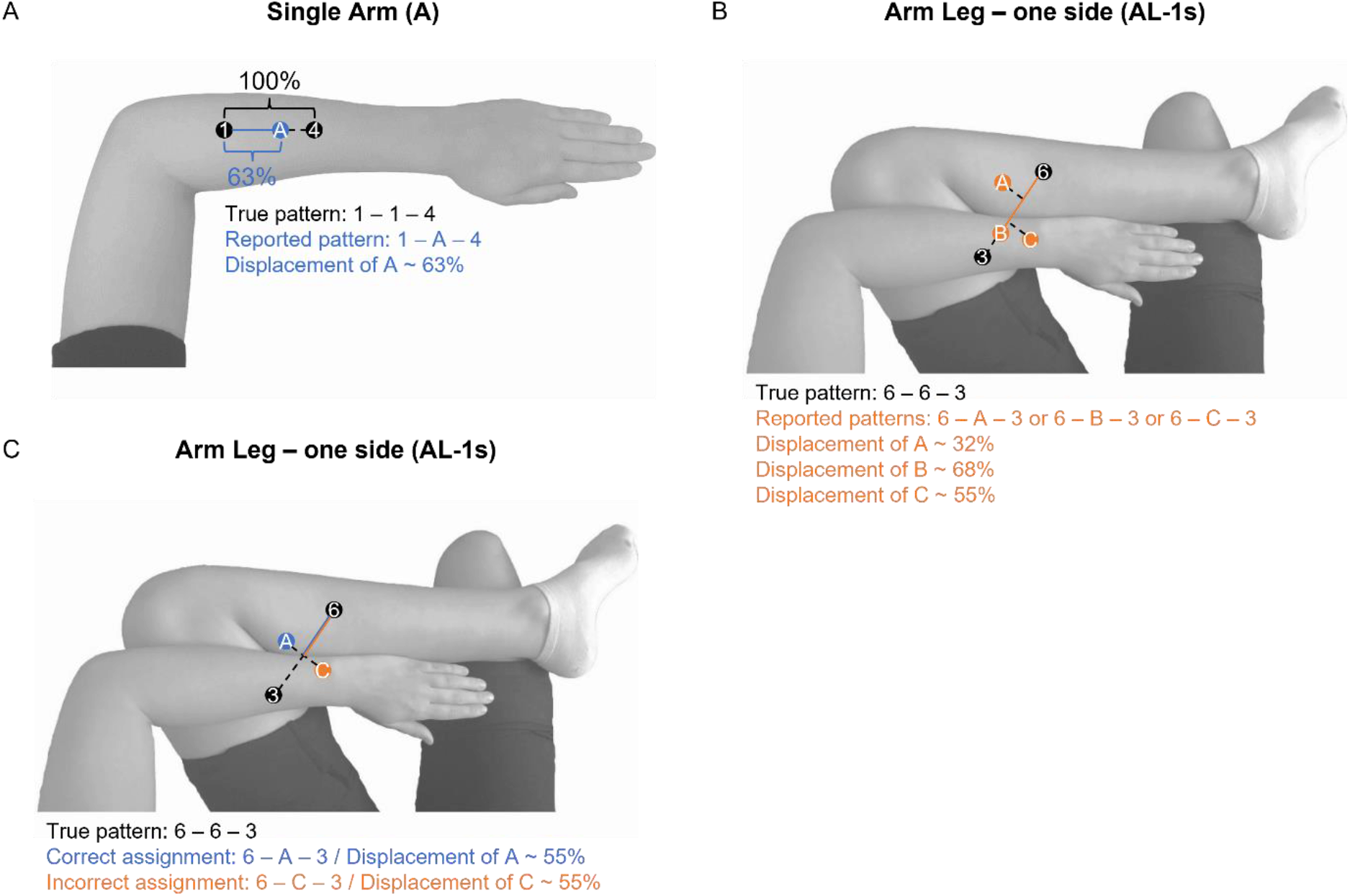
Schematic representation of the extracted measures, in single limb (A) and two-limb (B & C) configurations. The degree of displacement indicates how far the second stimulus was shifted towards the third. We considered the rabbit illusion as present when there was a displacement of the second stimulus > 10% of the distance between first and third stimuli. We further assessed whether the second stimulus was assigned on the correct limb, that is on the same as the first stimulus or not (C).

#### Statistical analysis

We used (Generalized) Linear Mixed Models ([G]LMM) for all statistical analyses, implemented in R (v3.6.1; RStudio v1.1.442; R Core Team, 2018) with the package lme4 (Bates et al., 2015). For each model, we implemented the maximum random effects structure for which the respective model still converged and improved model fit (Barr et al., 2013). We used the package car (Fox & Weisberg, 2011) to obtain statistics for main effects and interactions based on Type III Wald chi-square tests; we report these values rather than lme4 output). We performed post-hoc tests and contrast analyses with the package emmeans (Russell, 2019), and adjusted the resulting *p-values* with the false discovery rate method for multiplicity adjustment. Degrees of freedom for post-hoc tests of GLMM are infinite because they are calculated via the default asymptotic method to compare odds ratio in emmeans.

#### Statistical analysis: single stimuli

Aside from familiarizing participants with the stimuli, this condition tested whether participants discriminated between the different stimulated positions by testing for whether localization responses differed between all tactile locations. We ran a separate LMM for each limb configuration with factor *Stimulator* (numbers 1-6 and 1-8, respectively, for single and two-limb configurations), *Statistical analysis: three-stimulus sequences*. We performed separate comparisons of each control sequence type with the experimental illusory rabbit sequences, because the two control sequence types stimulated different spatial locations and were, therefore, not directly comparable. The time control sequences were designed as a comparison in which the first and second stimuli were in the same true location as the illusory rabbit sequences, but the expected percept differed between the two sequence types due to stimulus timing. The location control sequences were designed as a comparison in which the second stimulus was truly at an intermediate position between the first and third stimuli, that is, at a location at which the second stimulus may be illusorily perceived in the illusory rabbit sequences.

Data inspection revealed a bimodal distribution of the reported distance between first and second stimulus. One mode was made up of trials with a distance near 0 (see Figure S2), indicating that the second stimulus had been veridically perceived near its true location, implying that no illusion had been elicited (see Figure 2); this was more frequent in control sequences, a result that validates the experimental design. The other mode comprised trials in which the second stimulus was reported at some distance from the first, as would be expected if the rabbit illusion had been perceived. We modeled the bimodal distribution with a two-part modelling approach (Sauzet et al., 2019).

With this analysis strategy, we categorized, in a first step, whether the illusion was present or absent in a given trial by classifying a trial as “displacement absent” when the reported displacement of the second stimulus was <10% and as “displacement present” otherwise. Step 1 analyzed the frequency with which a shift was reported for the second stimulus in the different sequence types, especially whether this effect was modulated when stimulation spanned two limbs. This analysis used GLMMs with a binomial link function with factors *Limb Configuration* (A, L, AA, AL-1s, AL-2s) and *Stimulus Sequence* (illusory rabbit / time control or illusory rabbit / location control).

In a second step, we analyzed the magnitude and limb assignment of only those trials that had been classified as “displacement present” in the first analysis step. Thus, Step 2 assessed aspects of behavior only if the rabbit illusion had occurred. We first assessed the degree of illusory displacement of the second stimulus as an indicator of the rabbit illusion’s magnitude and analyzed whether localization of the middle stimulus differed between the three-stimulus sequences. We used an LMM with factors *Limb Configuration* and *Stimulus Sequence* (illusory rabbit vs. time control). We did not compare the illusory rabbit and location control sequences because the latter would render average reports of truly stimulated positions, and it would not be informative to compare those to the illusory rabbit sequences. Second, we assessed whether the second stimulus had been reported on the limb on which it had been presented, or whether participants reported this stimulus on the other, second limb. This effect could only occur in two-limb configurations, so that only the respective data entered this analysis. We used GLMMs with a binomial link function with factors *Limb Configuration* (AA, AL-1s, AL-2s) and *Stimulus Sequence* (illusory rabbit / time control or illusory rabbit / location control). By design, one would expect a second-stimulus displacement in many or all illusory rabbit sequence trials. Yet, as with most illusions, some participants will experience the illusion strongly and in almost all trials, and others will only experience it occasionally. In turn, by design one would expect no displacement in the time control sequences, because the long second SOA of 700 ms is supposed to eliminate illusory shifts of the second stimulus. However, long SOAs reduce, but do not fully eliminate, the occurrence of illusory displacements (Asai & Kanayama, 2012; Trojan et al., 2010; Warren et al., 2010). In our experiment, the number of illusory displacements varied considerably between participants.

For the above reasons, participants often reported a displacement in only very few trials in some condition. In such cases, averaging is likely misleading. Therefore, we excluded data points for which 5 or fewer trials were available for a given limb configuration and sequence type (distance displacement: 39 of 178 removed; limb identification errors: 29 of 159 removed). If we included all trials, the overall result pattern was qualitatively unchanged, but, unsurprisingly, there were a number of outliers suggestive of invalid interpretations.

### Experiment 2

#### Participants

We recruited 21 new participants for Experiment 2, of which one was excluded due to wrong placement of the stimulators. The remaining 20 participants (18 females, mean age=23 years, SD = 2.9) were all right-handed according to the EHI. Recruitment, inclusion criteria, handedness scoring, compensation, and laboratory setting were identical to those of Experiment 1.

#### General Procedures

The stimulation setup was identical to the left-arm-left-leg (AL-1s) configuration in Experiment 1, with identical positioning of the stimulators on the limbs (see Figure 2B). We restricted the experiment to stimulation of the left arm and leg, as this configuration had shown the highest proportion of reported stimulus displacement and stimulus assignment to the incorrect, second limb in Experiment 1. Participants performed a two-alternative force choice task (2AFC). They gave their responses with a two-button response box, which they operated with two fingers of their dominant hand. Participants wore headphones to cancel any noise from the stimulators and to receive feedback tones during the task. The session lasted for 1.5 to 2 hours. At the end of the session, participants judged stimulus intensities for arm and leg with a visual scale that ranged from zero (= no sensation) to ten (= very strong sensation). As in Experiment 1, participants rated stimulation stronger on the arm than on the leg (6.6 ± 2.1 vs. 4.2 ± 2.0; SE = .483, t = −4.99, p = .002).

#### Stimulus Sequences

Participants received tactile three-stimulus sequences with the same spatial locations and timings as in Experiment 1, that is, illusory rabbit, time control, and location control sequences For illusion and time control sequences, the first and second stimuli were presented on one limb, and the third on the other, just as in Experiment 1. Thus, the positions for the illusory rabbit and time control sequences were: 2-2-5, 3-3-6, 4-4-7, 5-5-2, 6-6-3, 7-7-4 (see Figure 2B). The positions for the location control sequences were: 3-5-6, 4-6-7, 5-3-2, 6-4-3, 2-3-5, 3-4-6, 6-5-3, 7-6-4. Each illusory rabbit pattern was repeated 50 times and each time control and location control pattern 25 times. In total, the experiment comprised 650 trials (300 illusory rabbit, 150 time control, and 200 location control trials). Sequence order was randomized and split into blocks of 50 trials. Participants took breaks after each block.

#### 2AFC task

Following the stimulus sequence, participants indicated by a button press on which limb they had felt *two* of the three stimuli Assignment of limb to response buttons was counterbalanced across participants. The task directly assessed whether the second stimulus was incorrectly assigned to the other limb, that is, whether it “jumped” from the truly stimulated limb to the other in the stimulus percept. As an example, think of a trial in which the first two stimuli were presented on the leg and the third on the arm. The first stimulus was always presented on the first limb, and the third stimulus on the second limb, and these stimuli had not been subject to substantial perceptual displacement in Experiment 1 (see also *Bayesian Modelling* section). Therefore, if the participant now indicated that (s)he perceived two stimuli on the arm, then (s)he must have perceived the second stimulus on the arm along with the third. Given that the second stimulus was always presented at the position of the first stimulus – here, the leg –, this response indicates that the second stimulus has been incorrectly localized to the arm. In other words, the second stimulus “jumped” from one limb to the other. Participants were instructed to press the correct button and respond correctly and as fast as possible after the three stimuli. The beginning of each trial was indicated by a beep (880 Hz, 100 ms). This was followed by a random interval of 100-500 ms, after which the stimulus sequence was presented. When participants did not respond within a 3000 ms time limit, measured relative to the offset of the last stimulus, a tone (440 Hz, 200 ms) reminded them to respond faster. Finally, a 1200 Hz tone feedback indicated that a response was considered invalid; this was the case if the response had occurred too fast (response time <100 ms after the sequence end; tone duration 100 ms) and if multiple button presses were registered (tone duration 200 ms). No-response and invalid trials were repeated at the end of a block.

After each trial, the program evaluated whether the response was correct and determined whether a confidence rating was shown. The inter trial interval was 500 ms if no confidence rating was presented.

#### Confidence ratings

We presented confidence ratings after some trials to assess how certain participants were about correct and incorrect choices. In Experiment 1, the rate of incorrect limb assignment was around 35%. Therefore, we increased the number of trials in Experiment 2 so that we would be able to reliably observe the phenomenon here as well. To keep the experiment’s duration low, we assessed confidence only after some randomly chosen trials. A confidence rating was shown with a probability of 80% after an incorrect response, that is, when the response indicated that the second stimulus had been assigned to the incorrect limb. After correct responses, a rating was shown with a probability of 20%. This seemingly unbalanced sampling was chosen to assess similar amounts of ratings for correct and incorrect choices, with incorrect choices occurring less frequently. Participants were not informed about whether their responses were correct or not and were unaware of our sampling strategy.

Confidence ratings had two phases. The participant was first asked, via text on the computer screen, whether the last button press had been correct. Participants responded yes/no by clicking on one of two buttons presented on the screen using the mouse. Following this binary question, a confidence rating asked how certain the participant was about the limb assignment. Participants adjusted a continuous slider with the mouse on the screen that indicated a line with 0, 50 and 100% landmarks. 100% meant that the participant was absolutely certain of their response; 0% meant that they were maximally uncertain. The currently selected value was continually shown under the rating line, and participants confirmed their choice by clicking on that number. Both yes/no and slider responses were unspeeded and did not have a time limit. After the confidence rating was completed, an inter-trial interval of 2000 ms gave participants time to remove the hand from the mouse and back on the response buttons before the next stimuli occurred.

#### Statistical analysis

As in Experiment 1, we ran separate analyses to compare illusory rabbit vs. time control and illusory rabbit vs. location control sequences. We devised three analyses. First, we assessed the probability of an incorrect response – indicative of a stimulus reported on the wrong limb – with GLMMs with a binomial link function with factor *Stimulus Sequence* (illusory rabbit / time control and illusory rabbit / location control).

Second, we analyzed only those trials in which participants had indicated that they thought to have been correct (94% of the trials with a confidence rating). We examined confidence ratings with a LMM with factors *Stimulus Sequence* (illusory rabbit / time control or illusory rabbit / location control) and *Accuracy* (correct, incorrect).

Finally, we investigated whether accuracy in Experiment 2 could be explained solely by confidence ratings. The underlying logic of this analysis was that localization errors would simply express uncertainty and, possibly, guessing, if differences across conditions are accounted for by confidence. In contrast, if confidence cannot explain our experimental data, then experimental effects must originate from other factors than uncertainty. We ran a GLMM with a binomial link function with factor *Stimulus Sequence* and a continuous covariate *Ratings*. The proportion of errors in control trials was quite low; therefore, we excluded the interaction between the two factors. Analysis tools and analysis strategies were otherwise identical to those reported for Experiment 1.

### Bayesian observer model

#### Application of the model for three stimuli

We adopted the Bayesian observer model by Goldreich and Tong (2013). The general logic of the Bayesian observer model for two stimuli is that the observer infers a movement trajectory (*x*_1_, *x*_2_) based on information about sensed positions (*x*_1*m*_, *x*_2*m*_), the time (*t*) that passed between them, the spatial uncertainty of each sensed position (σ_1_, σ_2_) and an assumption of low velocity (σ_*v*_) that expresses the observer’s expectation of location change in spatial units per one temporal unit. The low velocity prior reflects the average speed with which objects typically move along our skin in every-day life. Using Bayes’s theorem, the Bayesian observer model predicts the perceived positions computed as the mode of the posterior distribution, which is a two-dimensional Gaussian distribution with one dimension corresponding to *x*_1_ and the other to *x*_2_ when only two stimuli are involved.

In analogy, when the Bayesian observer model is extended to three stimuli, the resulting posterior distribution is a three-dimensional Gaussian. For three stimuli, the model integrates two time intervals: *t*_1_ as the interval between the first and second stimuli and *t*_2_ as the interval between the second and third stimuli. The posterior distribution is computed as described (Goldreich & Tong, 2013; see their Equation A49)

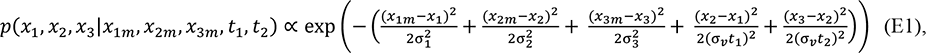

where *x*_*im*_ denotes the sensed positions of stimuli 1-3; σ_*i*_ their standard deviations, that is, the inverse of spatial precision, *t*_1_ and *t*_2_ the time intervals between stimuli (1, 2) and (2, 3), respectively; σ_*v*_ the low velocity prior; and *x*_*i*_ the true positions of the stimuli that the model attempts to infer given data and the prior assumption. The most probable configuration of *x*_*i*_ given the other parameters, is defined by the posterior mode and corresponds to the percept that the model predicts. The configuration of *x*_*i*_ corresponding to that mode can be computed analytically (for details see Goldreich & Tong, 2013 and our documented implementation on https://osf.io/a8bgt/).

#### Computation of model parameters

The model requires estimating the baseline localization variability for each stimulus as an estimate of σ_*i*_. We computed average localization as centroids and the standard deviations of localizations around the centroids for σ_*i*_ of each participant and limb configuration. We based centroids on responses during the three-stimulus sequences, rather than during single stimulus trials, because trial numbers for the latter were low and, thus, insufficient for obtaining reliable parameter estimates. For each stimulator, we averaged localization over all trials in which a sequence had begun with the stimulator of interest, because the first stimuli in our sequences were usually not subject to illusory displacement.

We obtained single, overall variability values for each limb in each limb configuration by first averaging standard deviation over x (horizontal/proximodistal) and y (vertical/mediolateral) directions for each stimulator and then across all stimulators. These per-limb variability estimates served as input to the model’s σ_*i*_ parameters.

Equations for the Bayesian Observer model are more straightforward for treating one than several dimensions. Therefore, we projected participants’ 2D centroids onto a straight line between the locations of stimuli 1 and 3. This procedure is the same as that for the attraction index we used as operationalization for illusory displacement (see section *Data processing*). Last, we transformed the data into approximate cm units. This step was necessary because the model’s low-velocity prior (σ_*v*_) is expressed as a value in cm/s in the literature (Goldreich, 2007; Goldreich & Tong, 2013) and is not transferable to the pixel units that we used for localization. We computed a transformation from pixel to cm units for each participant, based on the consistent 10 cm distance between the first and third stimuli throughout the entire experiment. We then transformed the projections for each limb configurations on the straight line into cm units by dividing the projected pixel values by the pixel value corresponding to 10 cm. This transformation was applied to both the *x*_*im*_ and the σ_*s*_.

Finally, we computed model predictions for each participant in all configurations and stimulus patterns. We used *t*_1_ and *t*_2_ parameters according to the stimulation intervals for the illusory rabbit and the time control sequences, respectively. For both sequences, we used a low-speed prior set to 10 cm/s, as suggested previously (Goldreich, 2007; Goldreich & Tong, 2013).

#### Statistical analysis

The predictions of the Bayesian observer model for the limb configurations depend on localization variability (inverse precision) of the stimulated limbs. We therefore compared the variability between the limbs prior to modeling. Because the number of levels of *Limb* varied between single-limb (only one limb) and two-limb configurations (two limbs), we analyzed the single-limb and the two-limb configurations in two separate models. We analyzed configurations A and L with an LMM with fixed factor *Limb Configuration*, and AA, AL-1s and AL-2s in an LMM with fixed factors *Limb Configuration* and *Limb* (limb 1 vs. limb 2 of the respective configuration).

We then assessed model performance in four ways. First, we compared the displacement of the second towards the third and of the third towards the first/second stimulus between the illusory rabbit and time control sequences using an LMM on the observed values including the fixed factors *Stimulus Sequence* (illusory rabbit vs. time control), *Limb Configuration*, and *Stimulus* (first, second, and third). Second, we focused on the model’s predictions about the displacement of the second stimulus on the leg compared to the arm in the illusory rabbit stimulus sequence. We compared the observed positions of the second stimulus in the illusory rabbit sequence for A and L with an LMM with fixed factor *Limb Configuration*, and AL-1s and AL-2s with an LMM with fixed factors *Limb Configuration* and *Direction* (stimulus sequence moving arm-to-leg vs. leg-to-arm).

Third, we assessed whether the model faithfully reproduces locations, and whether the model performs differently for single and two-limb configurations by regressing predicted onto observed positions. Participants typically mislocalized the second and third, but not the first stimulus. Including the first stimulus into model assessment would have artificially improved our measures of model performance; therefore we restricted analysis to the second and third. We tested for differences between single-limb and two-limb configurations by grouping single-and two-limb configurations into a factor *Limbs* (levels: one vs. two). The model’s predicted positions were entered as a continuous predictor.

Fourth, we analyzed whether the Bayesian model also predicted observed positions locally, i.e., whether it reproduces variation observed across participants when focusing on the variation within of the second and third stimuli, respectively. The model should be able to explain this variation because it is informed about the participants’ σ_*s*_. According to the model, participants with lower σ_*s*_ (higher precision) for a given stimulus position should display lower displacement of that position as compared to participants with lower precision. To test this prediction, we computed another LMM including the predicted values of the illusory rabbit pattern and the fixed factors *Limbs* and *Stimulus* (second vs. third).

## Results

### Experiment 1

#### Correct localization of single stimuli

In the single-location task, participants localized single stimuli to check for participants’ ability to differentiate and localize the different stimulus positions when no other stimuli are present that might trigger spatiotemporal integration (Trojan et al., 2010). Participants localized stimuli on the correct limb in 98.6% (AA), 99.3% (AL-1s) and 98.9% (AL-2s) of the two-limb configuration trials. There was a significant main effect of *Stimulator* for both one limb configuration (linear mixed models; A: χ²(5) = 1900; p < .001; L: χ²(5) = 1056; p < .001) as well as for each limb in two-limb configurations (AA – Arm 1: χ²(3) = 538; p < .001; AA – Arm 2: χ²(3) = 560; p < .001; AL-1s - Arm: χ²(3) = 713; p < .001; AL-1s - Leg: χ²(3) = 215; p < .001; AL-2s - Arm: χ²(3) = 379; p < .001; AL-2s - Leg: χ²(3) = 292; p < .001), indicating that participants’ localization responses differed significantly between the different stimulated locations. Post-hoc comparisons indicated that localization was indeed different for each true location (all p < .001, see Table S3). These behavioral differences suggest that participants perceptually discriminated between stimulated positions.

#### Higher proportion of illusory displacements in illusion than control trials

Having confirmed that participants were well able to localize the stimuli if they occurred alone, we inspected with the “three-location task” how stimuli affected each other in three-stimulus sequences.

In illusory rabbit sequences, 75.9% of participants’ second stimulus localizations were displaced towards the third stimulus by 10% or more of the distance between the first and third stimulus. Mislocalization by 10% distance or more was also evident in time control sequences, but at a lower proportion of 45.5%. Stimulus displacement in time control sequences, as seen here, is a regular finding (Asai & Kanayama, 2012; Warren et al., 2010), with a small displacement even at SOA of > 1000 ms between the second and third stimulus (Trojan et al., 2010). There are individual differences between participants regarding mislocalization in time control trials, and a number of participants indicated a 10% or higher displacement in only one or few trials (as shown by some of the very low frequencies in Figure 4). Importantly, the difference between the two sequence types was significant (GLMM, fixed main effect of factor *Stimulus Sequence,* χ²(1) = 39.61; p < .001), indicating a higher occurrence of illusory displacement in illusory rabbit than time control trials. There was also a significant main effect of *Limb Configuration* (χ²(4) = 93.7; p < .001) and an interaction between the two factors (χ²(4) = 35.23; p < .001). Nevertheless, post-hoc comparisons indicated that the probability to report a displaced second stimulus location was higher in illusory rabbit than in time control sequences for all limb configurations (all p < .001; see Figure 4 and Table 2). Moreover, the displacement occurred more often when all stimuli were presented on a single arm than in all other limb configurations (p < .021 for all post-hoc comparisons with the single arm (A) configuration; see the bigger difference between the means of the blue and red violins for this configuration in Figure 4). The remaining limb configurations did not differ from each other, suggesting that the proportion of reporting a displaced second stimulus was comparable across these limb configurations (all p > .327).

**Figure 4.**
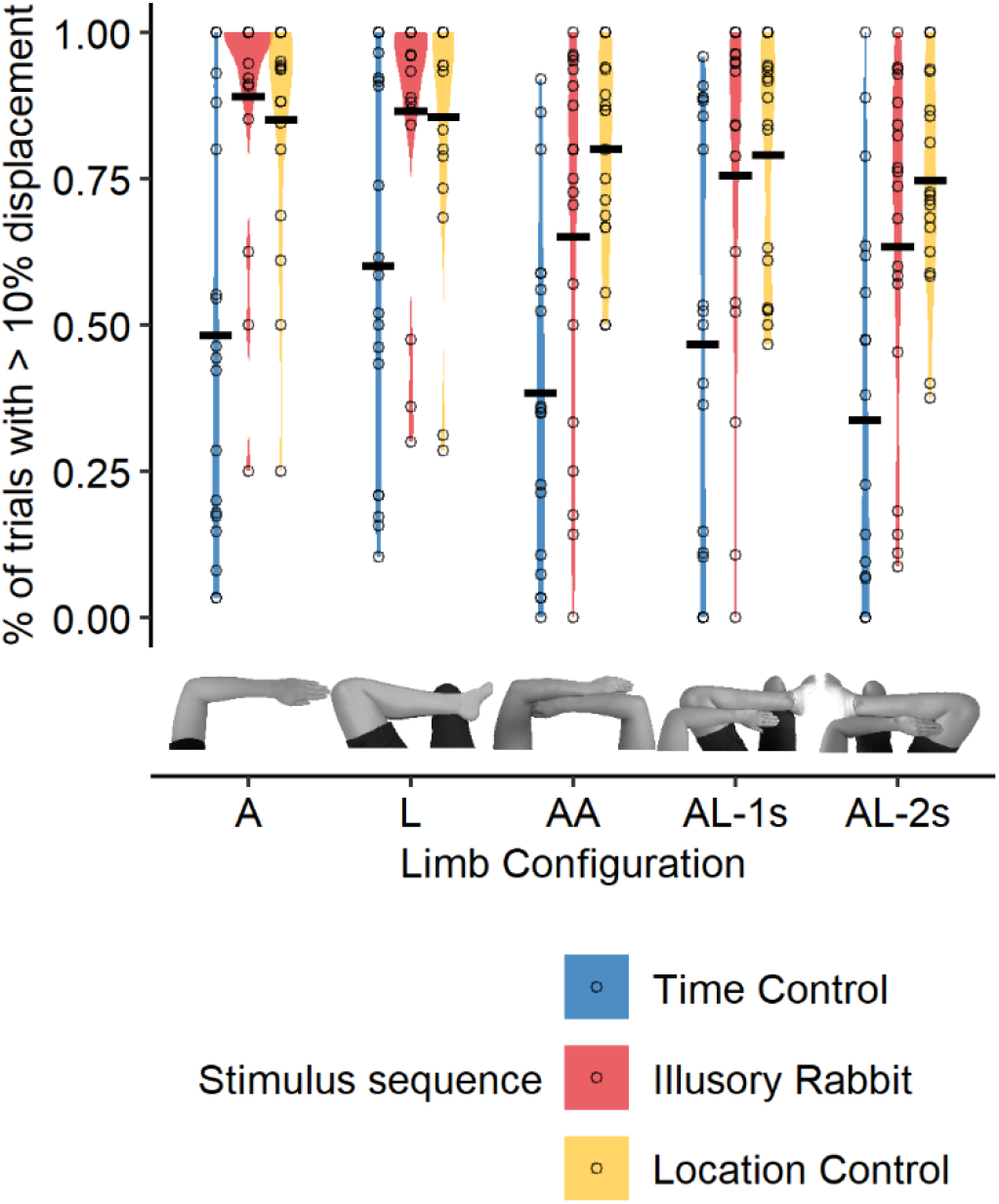
Frequency of reporting a displacement of the second stimulus of a sequence per Limb Configuration and Stimulus Sequence. Horizontal black line represents the mean frequency of reporting second stimulus as displaced from the first, and violins the distribution across all stimulus sequences of a given sequence type. Violins represent the distribution of the presented data, with thicker sections indicating higher density, that is, larger number of observations. Black dots represent the mean frequency of reporting a displaced second stimulus for single participants. A = Single Arm; L = Single Leg; AA = Two arms; AL-1s = Arm Leg – one side; AL-2s = Arm Leg – two sides.

**Table 2.** Post-hoc comparisons of the % of trials with a > 10% displacement of the second stimulus, for each stimulus sequence, in time control vs. illusory rabbit stimulus sequences. P values are FDR-corrected.

| stimulus sequence | limb configuration | odds ratio | SE | df | z ratio | p value |
| --- | --- | --- | --- | --- | --- | --- |
| Time control<br>vs.<br>Illusory Rabbit | Single Arm (A) | 0.044 | 0.01 | Inf | -10.42 | <0.001 |
|  | Single Leg (L) | 0.135 | 0.04 | Inf | -6.86 | <0.001 |
|  | Two arms (AA) | 0.203 | 0.06 | Inf | -5.80 | <0.001 |
|  | Arm Leg – one side (AL-1s) | 0.177 | 0.05 | Inf | -6.11 | <0.001 |
|  | Arm Leg – two sides (AL-2s) | 0.171 | 0.05 | Inf | -6.29 | <0.001 |

In sum, participants frequently reported a displaced second stimulus, indicative of having perceived the rabbit illusion, in the illusory rabbit sequences for all the limb configurations. Thus, the rabbit illusion occurs across two homologous limbs (AA) as well as across different types of limbs (AL-1s and AL-2s) and even across the body midline (AL-2s).

Not surprisingly, in location control sequences (three different stimulus locations; yellow violins in Figure 4), participants usually reported the second stimulus at a different location than the first – recall, that in these trials, the second stimulus was truly presented at an intermediate position. For comparability, we employed the same criteria to classify a difference in localization between two stimuli here as in the illusory rabbit sequences; in other words, we classified two stimuli to have been reported as spatially different if their locations differed more than 10% of the total reported sequence distance. Any deviation from 100% in this control condition, in which the stimuli were truly presented at different locations, can serve as an indicator for the upper bound of a reported displacement >10% in any other condition, for instance due to variability in indicating positions on the screen etc. With this criterion, the second stimulus was reported at a different location than the first in 80.9% of trials. This proportion is slightly, but significantly higher than in illusory rabbit sequences, as evident in a main effect of *Stimulus Sequence* (χ²(1) = 4.31; p = .038). There was no evidence for a modulation by the involved limbs, with neither a main effect of *Limb Configuration* (χ²(4) = 1.88; p = .758) nor an interaction between *Stimulus Sequence* and *Limb Configuration* (χ²(4) = 7.31; p = .121). Thus, although participants reported an illusory displacement in many illusory rabbit sequences, they did not do so in every single instance – on average, 5% less frequently than when a stimulus was truly presented along the trajectory. Critically, for time control sequences a displacement was reported, on average, 35% less frequently than for location control, and 30% less frequently than for illusory rabbit sequences. This comparison suggests that the percept evoked by illusory rabbit sequences is more similar to that of a truly displaced stimulus than that of a control sequence with a stimulus timing that discourages the rabbit illusion. Moreover, the small difference in reported displacement frequency suggests that the rabbit illusion emerges quite reliably, even if not with certainty.

#### Larger displacement in illusion than in control trials

Our previous analysis implies that illusory displacements were sometimes elicited in trials designed as non-illusion control trials, though with smaller frequency, as is usually reported in the literature (Asai & Kanayama, 2012; Trojan et al., 2010; Warren et al., 2010). This result may have two implications: either, the control sequences evoke the illusion with the same intensity, but only with lower probability; or they evoke a less intense illusion overall, resulting both in less frequent, but also smaller displacements. To assess this aspect, we restricted our analysis to trials in which a >10% displacement had been reported. Thus, displacement could range from a small, 10%, to a full, 100% displacement. Note, that the true distance between the first and third stimuli was identical in all the limb configurations and stimulus sequences. The displacement was 49% in illusory rabbit stimulus sequences, but only 34% in time control stimulus sequences (main effect of *Stimulus Sequence* (χ²(1) = 23.3; p < .001), indicating smaller illusory displacement in control than illusory rabbit sequences. Although there were a main effect of *Limb Configuration* (χ²(4) = 113.5; p < .001) and an interaction between the two factors (χ²(4) = 12.2; p = .016), post-hoc comparisons confirmed a significant effect of *Stimulus Sequence* for each limb configuration (all p < .001; see Figure 5 and Table 3). The displacement was smaller in the AA than in the AL-1s configuration (SE = 2. 4, t = −2.00, p = .023; see Figure 5). No other comparisons between limb configurations were significant (all p > .094; see *Table 4*).

**Figure 5.**
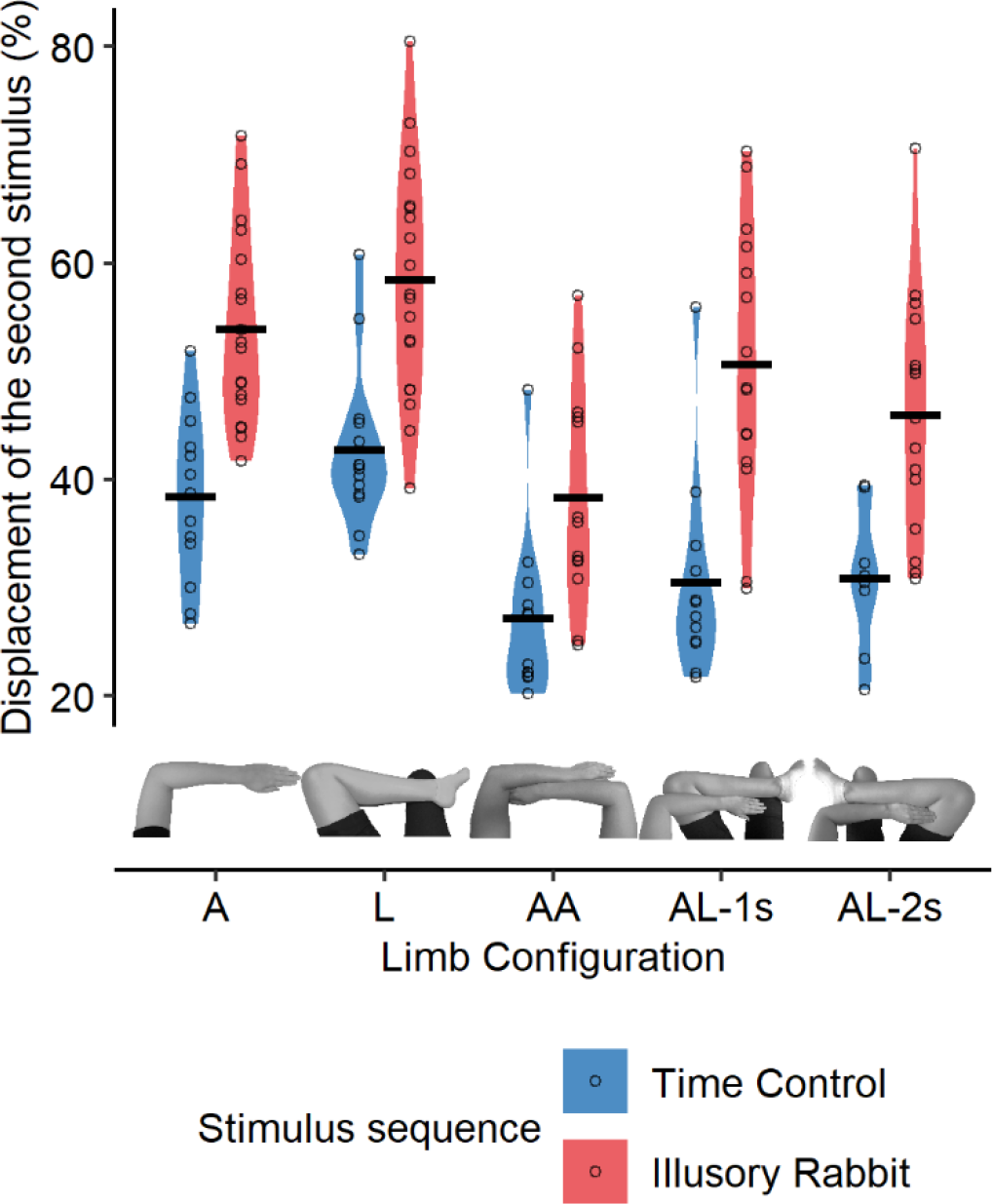
Displacement of the second stimulus, indicative of illusion strength, per Limb Configuration and Stimulus Sequence. Horizontal black line represents the mean displacement of the second stimulus, and violins the distribution across all stimulus sequences of a given sequence type. Violins represent the distribution of the presented data, with thicker sections indicating higher density, that is, larger number of observations. Black dots represent the mean displacement for single participants. A = Single Arm; L = Single Leg; AA = Two arms; AL-1s = Arm Leg – one side; AL-2s = Arm Leg – two sides. Data for the location control trials is not shown as it would be an average of truly stimulated positions, and it would not be informative to compare it to the illusory rabbit trials.

**Table 3.** Post-hoc comparisons of the degree of displacement, for each configuration, in time control and illusory rabbit sequences. Estimate indicates the differences in percent of displacement.

| stimulus sequence | limb configuration | estimate | SE | df | t ratio | p value |
| --- | --- | --- | --- | --- | --- | --- |
| Time control<br>vs.<br>Illusory Rabbit | Arm (A) | -14.3 | 2.3 | 29 | -6.20 | <0.001 |
|  | Leg (L) | -16.5 | 2.2 | 27 | -7.31 | <0.001 |
|  | Two arms (AA) | -10.3 | 2.6 | 50 | -3.88 | <0.001 |
|  | Arm Leg – one side (AL-1s) | -19.1 | 2.5 | 40 | -7.57 | <0.001 |
|  | Arm Leg – two sides (AL-2s) | -13.7 | 2.8 | 64 | -4.82 | <0.001 |

**Table 4.**
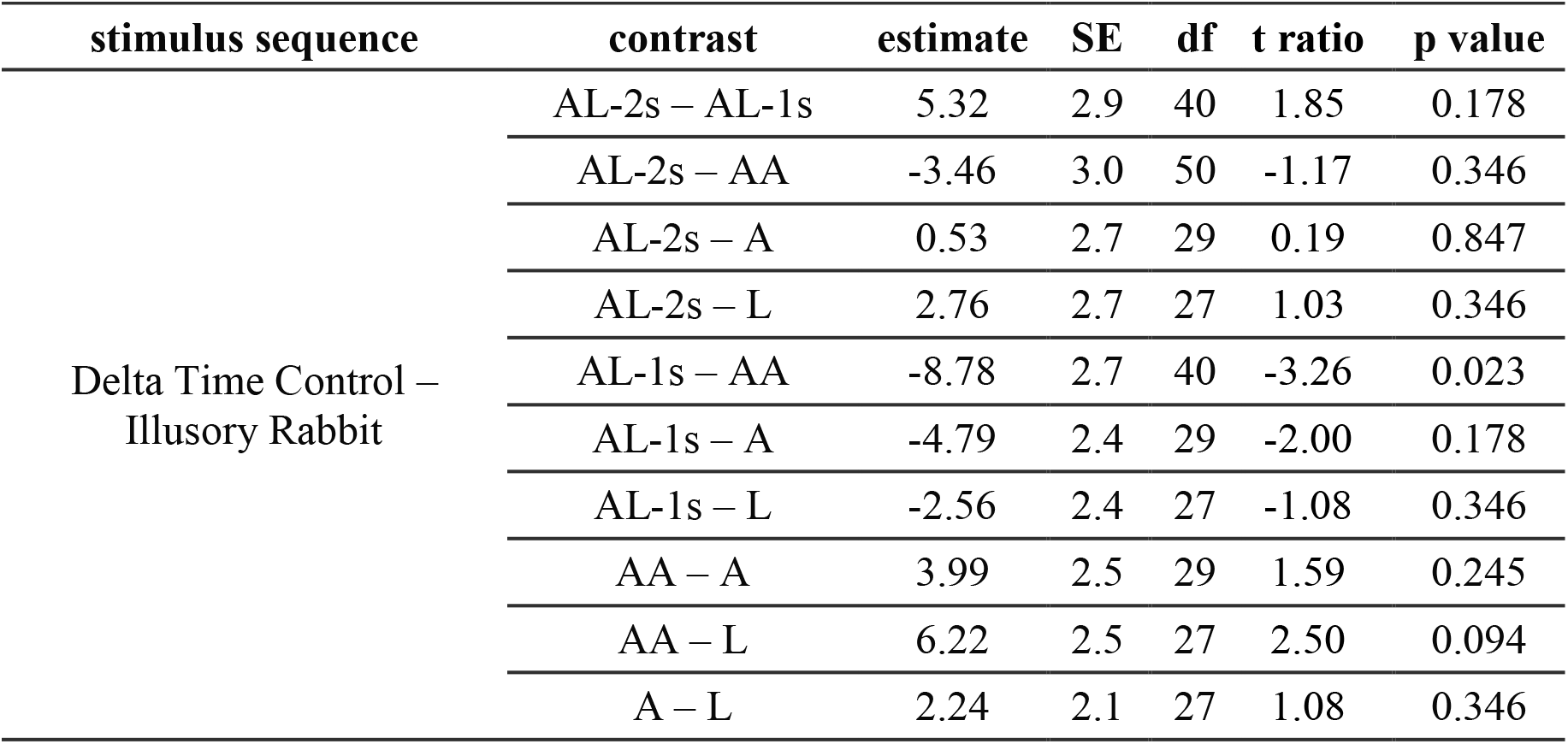
Post-hoc comparisons of the delta between Time Control and Illusory Rabbit sequences for each pair of configurations. Estimate indicates the differences in percent of displacement.

| stimulus sequence | contrast | estimate | SE | df | t ratio | p value |
| --- | --- | --- | --- | --- | --- | --- |
| Delta Time Control –<br>Illusory Rabbit | AL-2s – AL-1s | 5.32 | 2.9 | 40 | 1.85 | 0.178 |
|  | AL-2s – AA | -3.46 | 3.0 | 50 | -1.17 | 0.346 |
|  | AL-2s – A | 0.53 | 2.7 | 29 | 0.19 | 0.847 |
|  | AL-2s – L | 2.76 | 2.7 | 27 | 1.03 | 0.346 |
|  | AL-1s – AA | -8.78 | 2.7 | 40 | -3.26 | 0.023 |
|  | AL-1s – A | -4.79 | 2.4 | 29 | -2.00 | 0.178 |
|  | AL-1s – L | -2.56 | 2.4 | 27 | -1.08 | 0.346 |
|  | AA – A | 3.99 | 2.5 | 29 | 1.59 | 0.245 |
|  | AA – L | 6.22 | 2.5 | 27 | 2.50 | 0.094 |
|  | A – L | 2.24 | 2.1 | 27 | 1.08 | 0.346 |

In summary, these results indicate a larger displacement in illusory rabbit than in time control trials, both for single-limb and for two-limb configurations and irrespective of whether limbs belonged to the same body side or not.

#### The rabbit jumps from limb to limb

So far, we have shown that the displacement of the second stimulus is comparable when it involves homologous and non-homologous limbs, suggesting that it is not bound to skin space, but can arise in external space. The stimulus sequence for the illusion always presents the first two stimuli on one limb, and only the third stimulus on the other limb (e.g., Figure 2B, stimulus patterns 2-2-5, 6-6-3 etc.). Our sequences were arranged such that the distance between the first and third stimulus covered both involved limbs to an approximately equal amount. Therefore, if the illusion were based on interpolating a trajectory in 3D space, then the second stimulus should be perceived on the second limb when it was reported to be displaced by more than 50% towards the last stimulus. In such cases, the second stimulus would have been applied to one, but perceived on the other limb due to the rabbit-like interpolation of hops along the trajectory between the outer stimuli. As participants reported a large range of illusory locations for the second stimulus (see Figure 5), an assignment to the second limb should occur in some, but not all trials.

As previously, we restricted our analysis to trials in which the displacement of the second stimulus was > 10% and to sequences that stimulated both involved limbs (rather than just presenting all three stimuli to one of the two involved limbs). Participants reported the second stimulus on the wrong limb in 31.6% of illusory rabbit sequences. In contrast, this judgment error occurred in only 4.5% of time control sequences (main effect of *Stimulus Sequence* (χ²(1) = 17.7; p < .001). The main effect of *Limb Configuration* (χ²(2) = .88; p = .64) and the interaction between the two factors (χ²(2) = 3.43; p = .18; see Figure 6) were not significant. Thus, participants attributed the second stimulus to the incorrect limb particularly in illusory rabbit sequences, independently of which two limbs were stimulated.

**Figure 6.**
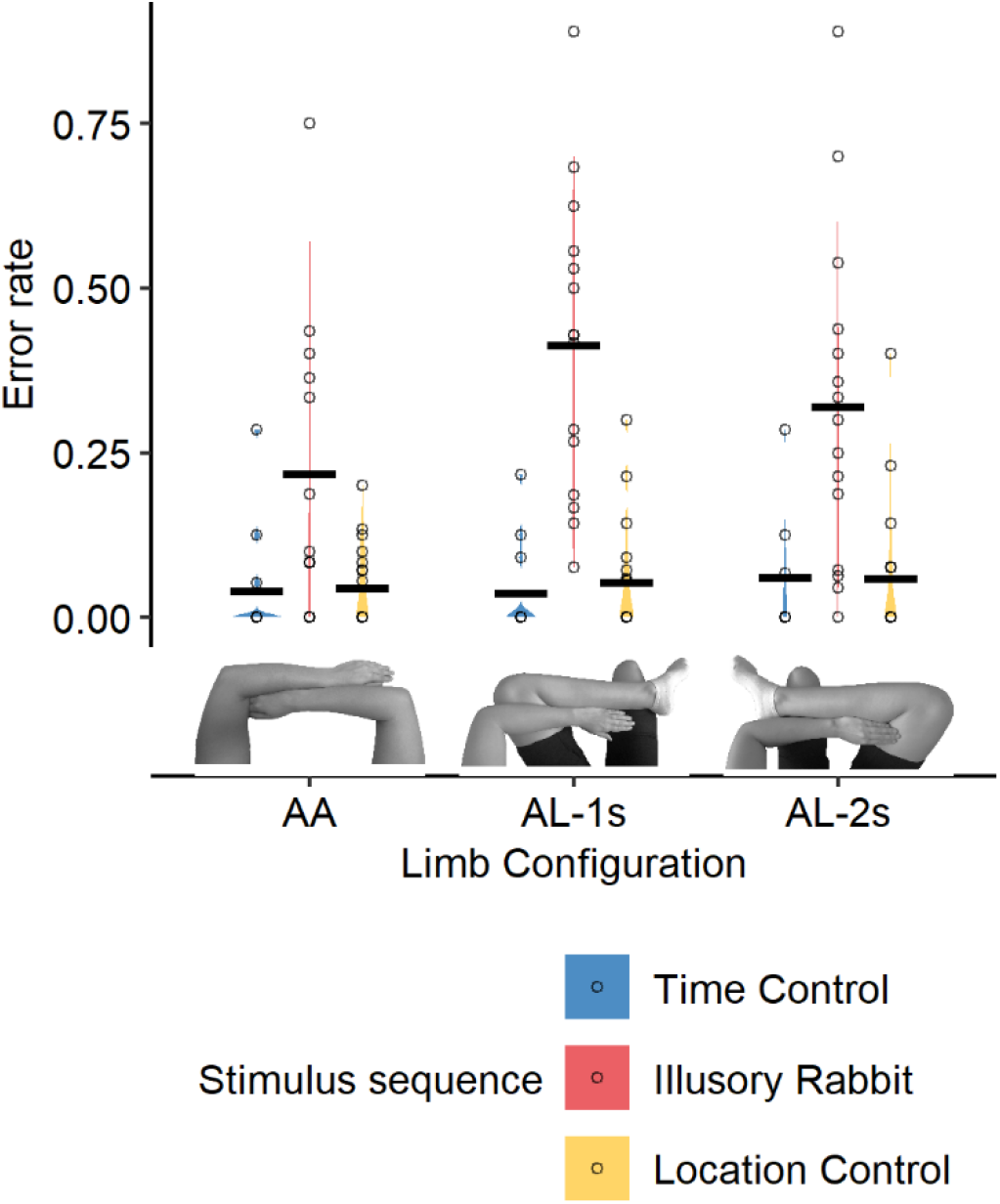
Error rate per Limb Configuration and Stimulus Sequence. Horizontal black line represents the mean error rate, and violins the distribution across all stimulus sequences of a given sequence type. Violins represent the distribution of the presented data, with thicker sections indicating higher density, that is, larger number of observations. Black dots represent the mean error rate for single participants. AA = Two arms; AL-1s = Arm Leg – one side; AL-2s = Arm Leg – two sides.

In our second type of control sequence, the location control sequences, stimuli occurred at three distinct locations. Here, the second stimulus occurred on the first limb in some, and on the second limb in other trials. Participants usually reported the correct limb (mean error: 5.2%), significantly different from the proportion of errors in illusory rabbit sequences (main effect of *Stimulus Sequence* (χ²(1) = 15.85; p < .001; see Figure 6). Again, there was no evidence for differences between limb combinations (main effect of *Limb Configuration* (χ²(2) = 2.08; p = .35; interaction of *Stimulus sequence* × *Limb Configuration*, χ²(2) = 4.53; p = .10; see Figure 6). Note, that errors were comparable in the two types of control sequences, and, thus, that frequent attribution to an incorrect limb was, therefore, a phenomenon unique to illusory rabbit sequences.

In sum, these results suggest that the second tactile stimulus perceptually jumped to the other limb in about one third of those trials in which the rabbit illusion was elicited. Incorrect assignment occurred in all two-limb configurations alike, including configurations that involved an arm and a leg. In other words, when the trial involved an arm and a leg, the middle stimulus was frequently mislocalized from an arm to a leg, or vice versa.

This finding entails that the rabbit’s hops are not just distributed along the skin, but that the rabbit can hop to a distant and non-homologous body part, presumably because the tactile stimulation pattern implies a movement in space towards the second limb. Thus, it suggests that the influence of expectation on perception is so strong that even the limb where touch is felt is “corrected”, a remarkable effect that defies explanations of the rabbit effect in terms of interactions between peripheral receptive fields or S1 neighboring neurons. Notably, previous studies have observed such jumps between body parts only between fingers of a same hand (Trojan et al., 2014; Warren et al., 2010), that is, skin regions that are represented in neighboring areas of primary somatosensory cortex of one hemisphere.

However, some alternative explanations need to be ruled out. One consideration is that, in our setup, participants reported the location of the three tactile stimuli on a trackpad, based on a photograph that displayed their current limb configuration. It is possible that participants did not always respond with high accuracy. Participants may have attempted to indicate the border region of the first limb, but accidentally indicated a region on the picture that already belonged to the other limb. However, visual inspection of the distribution of reported locations on the photographs did not suggest that this was the case: most responses were, in fact, clearly inside one or the other limb (see Figure S3).

Another consideration is that the fast stimulation we employed to elicit the rabbit illusion may have increased the level of uncertainty about the stimuli’s locations compared to the slower stimulation patterns of the control sequences. Accordingly, participants may have been unsure about where the second stimulus had occurred and may then have chosen a random location on the photographs without actually having had a clear percept of a stimulus on that limb.

We addressed these alternative explanations with a new experiment in which we explicitly assessed on which limb the second stimulus had been perceived and how confident participants were about their judgments.

### Experiment 2

#### Illusory jumps to the other limb occur irrespective of response mode

We presented the same three-stimulus sequences as in Experiment 1, but only to an arm and a leg (AL-1s). For each sequence, participants indicated whether two of the three stimuli had occurred on the first or on the second limb. In illusory rabbit and time control sequences, the first two stimuli are identical and, thus, always occurred on the first limb. The response that two stimuli had occurred on the second limb, therefore, indicates that the second stimulus was assigned to the incorrect limb, presumably because the rabbit illusion had emerged. Participants reported two stimuli on the second limb in 12.3% of the trials in illusory rabbit sequences, but only in 5.1% of time control sequences (χ²(1) = 37.2; p < .001; Figure 7). As for the location control sequences, participants made errors in 6.8% of the trials (χ²(1) = 9.5; p = .002; Figure 7), that is they sometimes reported two stimuli on the first limb when they were really occurring on both limbs (e.g. 3-5-6) and sometimes reported two stimuli on the second limb when they were really on the first one (e.g. 2-3-5).

**Figure 7.**
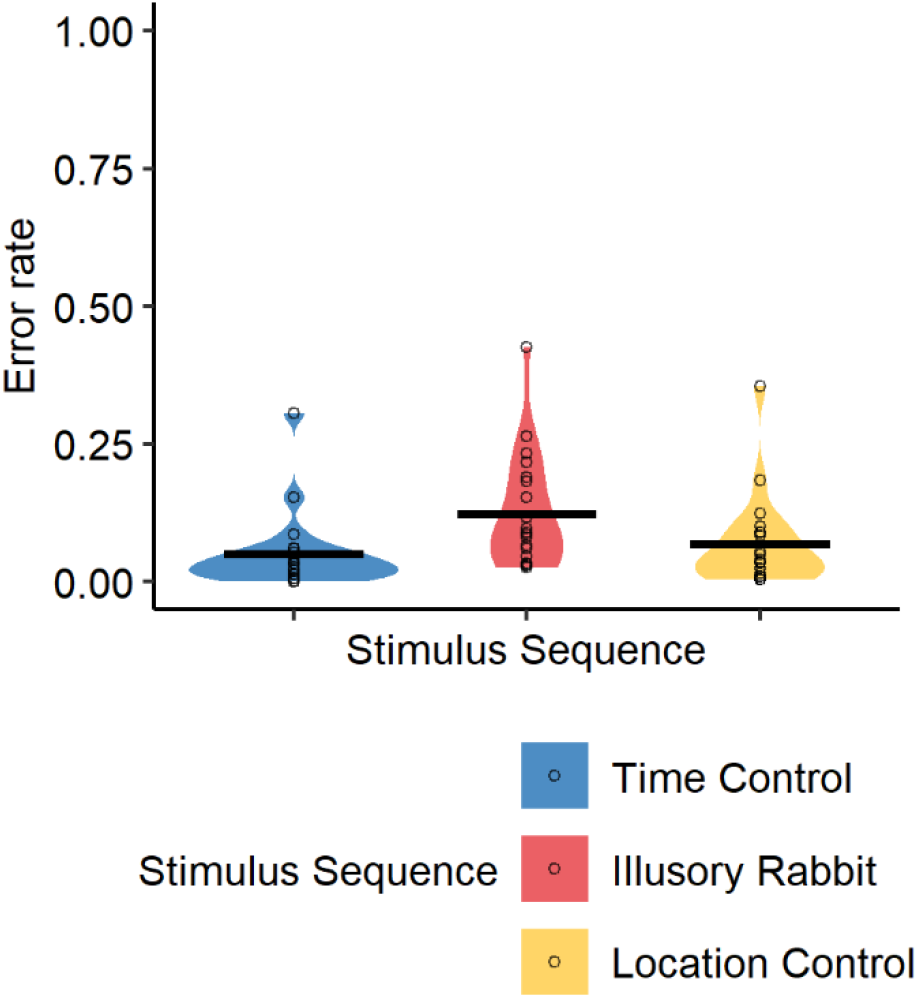
Error rate per Stimulus Sequence. Violins represent the distribution of the presented data, with thicker sections indicating higher density, that is, larger number of observations. Horizontal black line represents the mean. Black dots represent mean error for single participants.

Thus, Experiment 2 corroborates our finding from Experiment 1 that incorrect assignment can occur between arms and legs. Errors were less frequent in Experiment 2 (compare Figure 6 and Figure 7). When based on the total number of trials, an incorrect limb assignment occurred in 31.8% of Experiment 1’s illusory rabbit trials. This difference in numbers is likely due to methodological factors. First, Experiment 1 involved presenting participants with a photograph of their limb configuration; this visual information may have influenced the spatial judgments made by participants. Second, Experiment 2 posed only a single binary question directly following stimulation; in contrast, Experiment 1 required reporting all three stimuli, and the prolonged occupation with stimulus localization may have influenced participants’ responses. Importantly, the lower frequency of errors across limbs does not mean that the rabbit illusion occurred less frequently in Experiment 2. The measure was designed to exclusively assess incorrect assignment across limbs but does not distinguish whether the second stimulus was perceived as displaced but nonetheless correctly perceived on the first limb, or whether no displacement was perceived at all.

#### Rabbit jumps are not a result of perceptual uncertainty

Having replicated that participants wrongly assign stimuli from one limb to the other, we next turned to the question whether such reports may have simply resulted from confusion and uncertainty evoked by the unusual stimulation pattern. To this end, we compared reported confidence about the responses across sequences. We excluded trials in which participants stated that they had responded incorrectly (5.2% of ratings). Given that we obtained ratings in ∼80% of trials with an incorrect response without participants being aware of an incorrect answer, our data contained many trials in which participants reported a stimulus on the wrong limb but were obviously unaware of their error.

As before, we compared illusory rabbit sequences separately to time control and location control sequences. Both LMMs revealed a significant interaction between *Accuracy* (correct/incorrect) and *Stimulus Sequence* (illusory rabbit vs. time control: χ²(1) = 12.7; p < .001; illusory rabbit vs. location control: χ²(1) = 4.60; p = .032). Confidence was comparable across the different stimulus sequence types when participants gave incorrect responses (illusory rabbit sequences: 66.7%; time control sequences: 69.3%; location control sequences: 72.9%; all post-hoc comparisons are detailed in Table S4; see Figure 8). Recall that errors in illusory rabbit trials indicate that the second stimulus was perceived on the wrong limb; thus, there is no evidence that such mislocalization was accompanied by a reduction in confidence compared to any other stimulus sequence.

**Figure 8.**
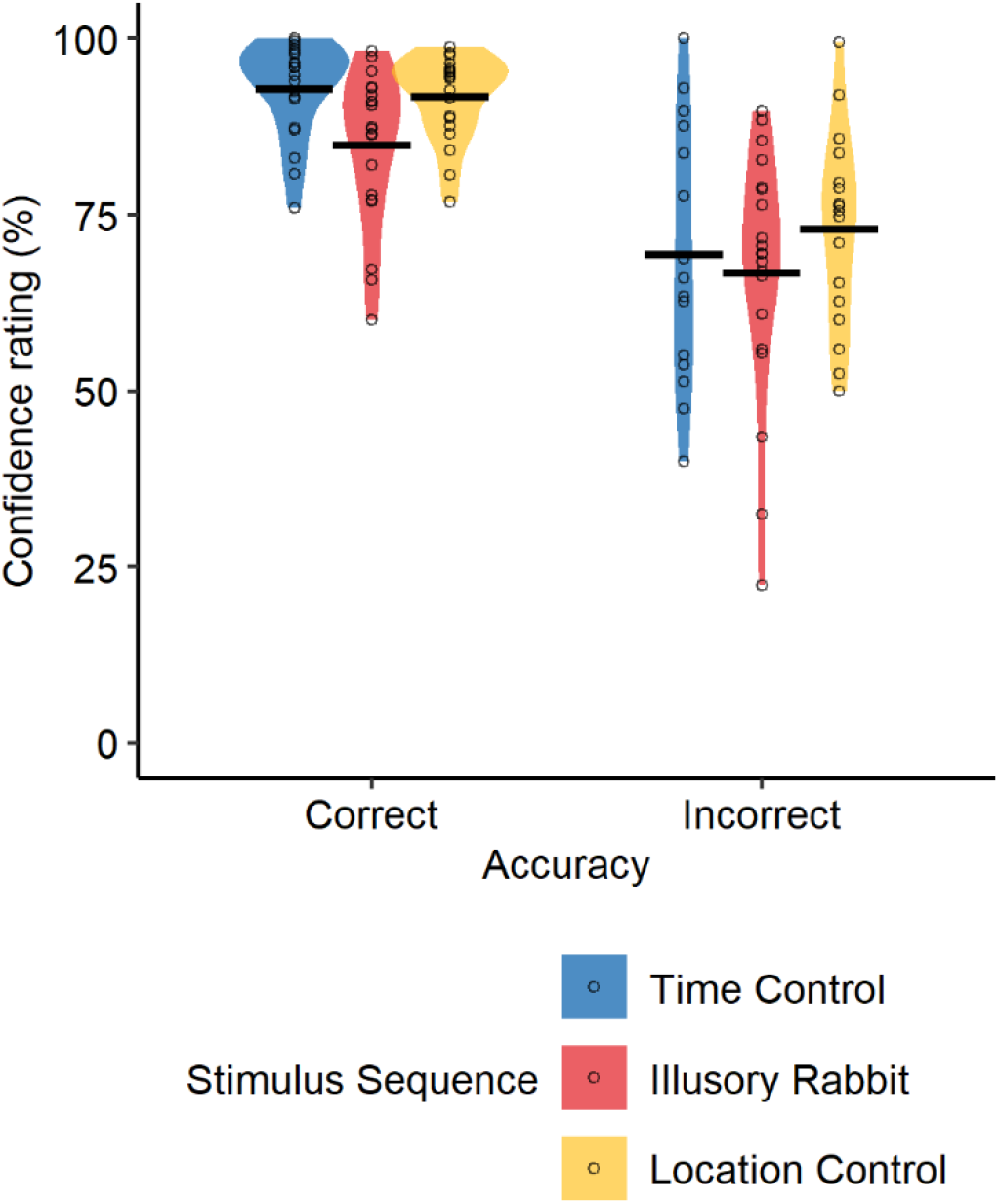
Confidence ratings in Experiment 2 for the different stimulus sequences, when participants were correct vs. incorrect. Incorrect response in time control and illusory rabbit sequences imply that the second stimulus was perceived on the wrong limb. Violins represent the distribution of the presented data, with thicker sections indicating higher density, that is, larger number of observations. Horizontal black line represents the mean. Black dots represent mean error for single participants.

As is common, confidence was higher for correct than for incorrect responses (e.g. Badde et al., 2019; Fleming & Frith, 2014), indicated by significant main effects of *Accuracy (*χ²(1) = 63.8; p < .001 and χ²(1) = 52.0; p < .001 for illusory rabbit vs. time control and location control respectively). However, when participants gave a correct response, their confidence was lower in illusory rabbit than in time control sequences (84.8% vs. 92.8%; t(21) = 5.8, p < .001) and location control sequences (84.8% vs. 91.7% respectively; t(23) = −4.2, p < .001; see Figure 8). Given that the number of incorrect responses differed across sequence types, but confidence was comparable for incorrect responses of all sequences, this suggests that uncertainty is unlikely to have played a role in the emergence of the rabbit illusion. Yet, we further tested whether uncertainty fully explains sequence differences in our experiment by adding confidence ratings as a covariate in a GLMM. If errors were merely a by-product of uncertainty, then any differences between illusory rabbit and time control sequences should be eliminated once variance induced by confidence ratings is accounted for. To the contrary, the main effect of *Stimulus Sequence* was significant also when *Confidence Rating* was included as covariate (χ²(1) = 30.1; p < .001).

In sum, differences in perceptual reports across stimulus sequences were not explained by differences in confidence. Thus, incorrect assignment of the second stimulus from one limb to another in the context of the rabbit illusion was not caused by perceptual uncertainty.

### Bayesian observer model

We observed a rabbit effect between two limbs, suggesting that the cutaneous rabbit illusion is based on external, not only anatomical space. But is this a reflection of statistically optimal integration as suggested by Bayesian accounts of perception? A Bayesian observer model (Goldreich, 2007; Goldreich & Tong, 2013) has been proposed as a framework for the cutaneous rabbit effect but has so far only been applied to single-limb scenarios where anatomical and external space cannot be dissociated. Many events involve straight or near-straight motion in space; accordingly, we expect that the brain’s priors should be optimally tuned to near-straight motion trajectories in external space. This view implies that the Bayesian observer model should capture participants’ behavior when we consider the Euclidian distance between the locations of our stimulus sequences. In particular, distance on the skin is large for all two-limb configurations of our experiments, whereas the Euclidian distance was comparable across all configurations. Therefore, we expect the Bayesian observer model to accurately account for the rabbit effect in all limb configurations when we feed it with external-spatial, that is, Euclidian, distance. On the one hand, this outcome would support that space and velocity, as conceptualized in the Bayesian model, are externally based. On the other hand, it would indicate that spatiotemporal integration based on priors is a feature grounded in 3D interaction space and, thus, is not restricted to anatomical space or processing within a single hemisphere. The model predicts where participants perceive each stimulus of a tactile sequence based on statistically optimal integration of sensory input with a “low velocity prior”. This prior expresses the brain’s experience-based prediction that tactile stimuli which follow each other close in time are usually close together in space and regularly originate from one common event or object that moves comparably slowly. For stimuli that imply fast motion, as in the cutaneous rabbit paradigm, the prior shapes perception in a way that stimuli are perceived as closer to each other than they really were, resulting in the observed shift of the second stimulus towards the third. However, the influence of the prior depends on the spatial variability of the stimulus: when variability is high – that is, when the true stimulus’s position of the current sensory input is uncertain – then prior experience will be given more weight, effectively “correcting” sensory input towards what is deemed plausible based on experience.

#### Higher variability for tactile localization on the leg than on the arm

Previous research has shown that tactile localization variability strongly varies between different body parts and is lower on the leg than on the arm (Mancini et al., 2014; Weinstein, 1968). In the limb configurations of Experiment 1, the Bayesian observer model, therefore, predicts a stronger rabbit effect on the leg than on the arm. Moreover, for the arm-leg limb configurations it predicts that the rabbit effect depends on direction. It should be stronger when the first two stimuli are presented on the leg, where variability is higher, compared to when they are presented on the arm, where variability is lower. With respect to apparent motion speed, the differences in participants’ reports evoked by our illusory rabbit vs. time control sequences illustrate that the rabbit effect depends on stimulus timing: stimulus timing is faster in illusory rabbit than in time control sequences, so that the Bayesian model predicts greater rabbit effects in the former, consistent with our findings.

We computed localization variability for the different limbs involved in our Experiment 1 and then fitted the Bayesian observer model to single-participant data for illusory rabbit and time control sequences in the different limb configurations. We then compared predicted with participants’ observed behavior.

Variability was higher, and thus, precision lower, on the leg than on the arm (see Figure 9). This was evident from a significant main effect of *Limb Configuration* when comparing the single arm and leg configurations in an LMM (χ²(1) = 22.2; p < .001). Comparison of AA, AL-1s and AL-2s configurations in a second LMM that included *Limb* and *Limb Configuration* as factors revealed a significant interaction between the two factors (χ²(2) = 91.4; p < .001); post-hoc analyses further showed a limb difference for AL-1s and AL-2s, each with higher variability for the leg. In contrast, no difference was evident between the two arms in AA (see Table S5 for pairwise comparisons).

**Figure 9.**
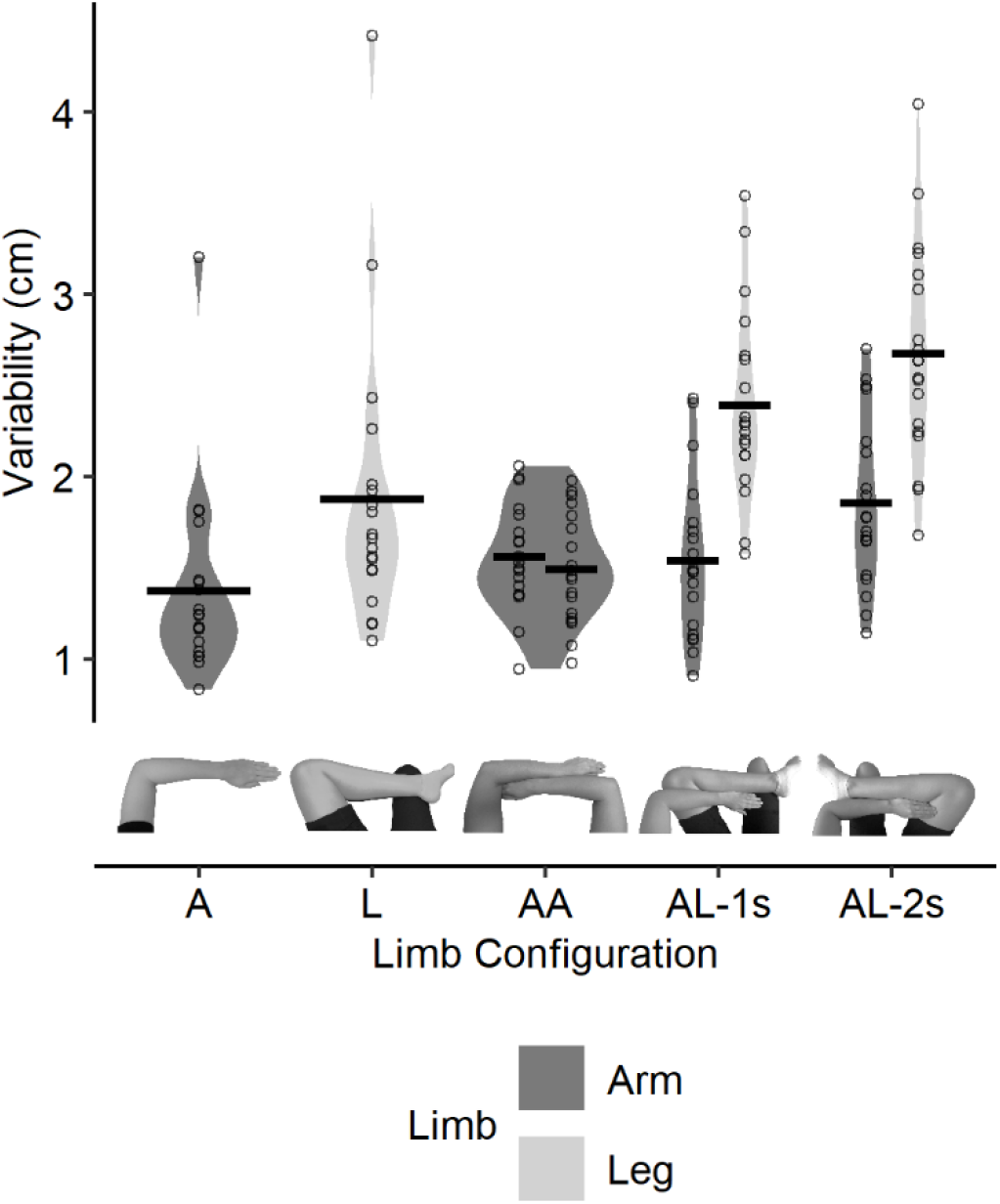
Variability of participants’ localization performance for the different limb configurations. Each point represents the averaged variability (x and y direction combined) for each participant. Lower variability indicates higher precision. Note that values were generally higher on the leg than on the arm, indicating lower tactile precision on the leg. A = Single Arm; L = Single Leg; AA = Two arms; AL-1s = Arm Leg - one side; AL-2s = Arm Leg - two sides.

#### Applying the model to three-stimulus sequences

We evaluated four predictions of the Bayesian Observer Model (see Figure 10A). First, participants should localize the first stimulus in both illusory rabbit and time control stimulus sequences with high accuracy, which is evident from the predicted locations being close to the zero line that represents the veridical position of the first and second stimuli. This prediction is due to the long SOA of 700 ms between the first and second stimulus; this long time interval diminishes the influence of later stimuli on the first one.

**Figure 10.**
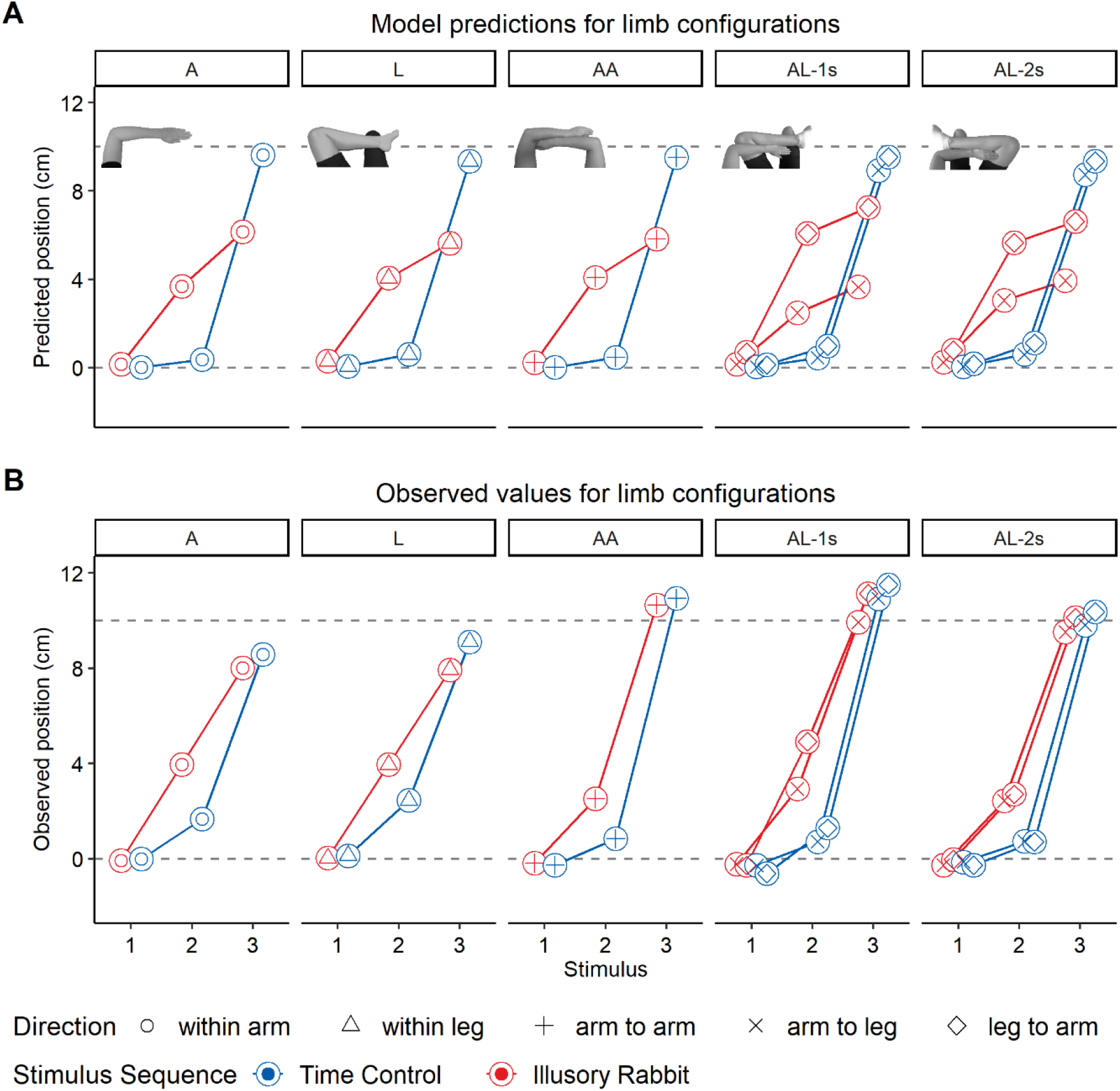
Predicted (A) vs. observed positions (B) for the limb configurations in illusory rabbit and time control sequences. Positions of the stimuli were predicted using an extended Bayesian observer model based on the model described by Goldreich and Tong (2013). The points represent mean positions (averaged over participants) in cm units along a straight line between the centroids of the respective stimuli. The dashed horizontal line at the y-value of zero represents the veridical position of the first and second stimuli and the dashed line at the y-value of 10 the veridical position of the third stimulus for both stimulus sequences. Illusory displacement is indicated by the second (and in some instances the third) stimuli falling in the area between the two dashed lines. A = Single Arm; L = Single Leg; AA = Two arms; AL-1s = Arm Leg - one side; AL-2s = Arm Leg – two sides.

Second, for illusory rabbit sequences, the model predicted strong attraction of the second towards the third stimulus and vice versa, due to the short time interval between them. For time control sequences, localization for the second and third stimuli should be close to their veridical positions.

Third, the model predicted slightly stronger displacement of both second and third stimuli in the single leg (L) compared to the single arm (A) configuration, given the higher variability on the leg than on the arm (Figure 9).

Fourth, the model predicted direction-dependent illusory displacement for AL-1s and Al-2s, again based on the differences in variability between arms and legs: For AL-1s and Al-2s, the displacements were predicted to be stronger for the second stimulus as well as weaker of the third stimulus for trials with a leg-to-arm than for trials with an arm-to-leg direction.

#### Similar patterns for predicted and observed displacement

The experimentally observed positions are shown in Figure 10B. The observed behavior patterns largely resemble those predicted by the model (see Figure 10A). This is most evident from larger displacement values for the second stimulus for the illusory rabbit compared to the time control sequences in all limb configurations (difference = 1.97 cm; t(4602) = 17.5, p < .001). Across all limb configurations, there was also a significantly stronger displacement of the third towards the first/second stimulus in the illusory rabbit compared to the time control sequences (difference = 0.69 cm; t(4602) = 6.2, p < .001). These results confirm the attraction of the second and third stimuli towards each other in the illusory rabbit sequences. In line with the model, the position of the first stimulus did not differ significantly between the stimulus sequences (difference = 0.01 cm; t(4602) = 0.1, p = .903).

Observed behavior did not match the model’s prediction of higher perceived displacement for single leg (L) over arm (A) sequences (χ²(1) = 0.51; p = .48). In contrast, the direction-dependent displacement patterns predicted for AL-1s and Al-2s were partially reflected in the observed behavior: the displacements of the second stimulus in the illusory rabbit sequences were significantly higher for the leg-to-arm than for the arm-to-leg direction in the AL-1s (difference = 1.96 cm; t(44) = 3.28, p = .004), but not in the AL-2s (difference = 0.02 cm; t(44) = 0.04, p = .966). The difference in the direction-based effect between these conditions was significant, indicated by the interaction between *Limb configuration* and *Direction* (χ²(1) = 5.5; p =.019). These results suggest that localization variability of individual limbs affect perceptual reports in the two-limb configurations, in line with a proposed process of Bayesian integration across limbs and, moreover, that this process more strongly applies when arms and legs belong to the same body side.

Our model-data comparisons have so far addressed specific hypotheses derived from the Bayesian model; next we evaluated its overall fit to the data when combining all available data in a mixed-effects regression analysis that predicts the observed with the predicted positions. The aim was to evaluate whether there are any differences in model fit between the single-limb configurations, for which the model was originally conceptualized, and the two-limb configurations for which the model has not yet been evaluated. We compared the AA, AL-1s, and AL-2s limb configurations against the single limb (A, L) conditions (analyses for each individual limb configuration are shown in Table S6). In a first step, we computed an LMM that predicted participants’ localization responses from the model’s estimates across single- and two-limb configurations. If the model correctly reflected behavior, then plotting observed against predicted localization values would fall onto a unity line of slope 1. The LMM revealed a significant linear relationship between observed and modelled localization responses for both single-limb (b = .80, t(2183) = 27.9, p < .001) and two-limb (b = 1.1, t(2183) = 47.5, p < .001) limb configurations. However, slopes differed significantly between the two (χ²(1) = 82.2; p < .001; see Figure 11). For the single-limb configurations, the slope was < 1, driven largely by the larger-than-predicted displacements of the second stimulus in the time control sequence (see Experiment 1 and discussion there). This displacement in the control sequence also deviates from the predictions of the Bayesian observer model, suggesting that other factors beyond weighted integration play a role in the perception of tactile location. For the two-limb configurations, the slope was > 1, driven mainly by the smaller-than-predicted displacements of the third stimulus, especially in the illusory rabbit sequence. This discrepancy is probably the most obvious mismatch between the Bayesian model and our experimental data. In Figure 11, this mismatch is evident by the third stimulus (red circle) being far above the diagonal line. This location indicates that participants reported the third stimulus closer to its veridical position than the model predicted. An additional LMM revealed that this discrepancy was significant in single-limb (difference = 2.1 cm; t(32.4) = 8.46, p < .001) and two-limb configurations (difference = 4.8 cm; t(24.6) = 20.85, p < .001), but significantly larger in the two-limb configurations (χ²(1) = 162.3; p < .001).

**Figure 11.**
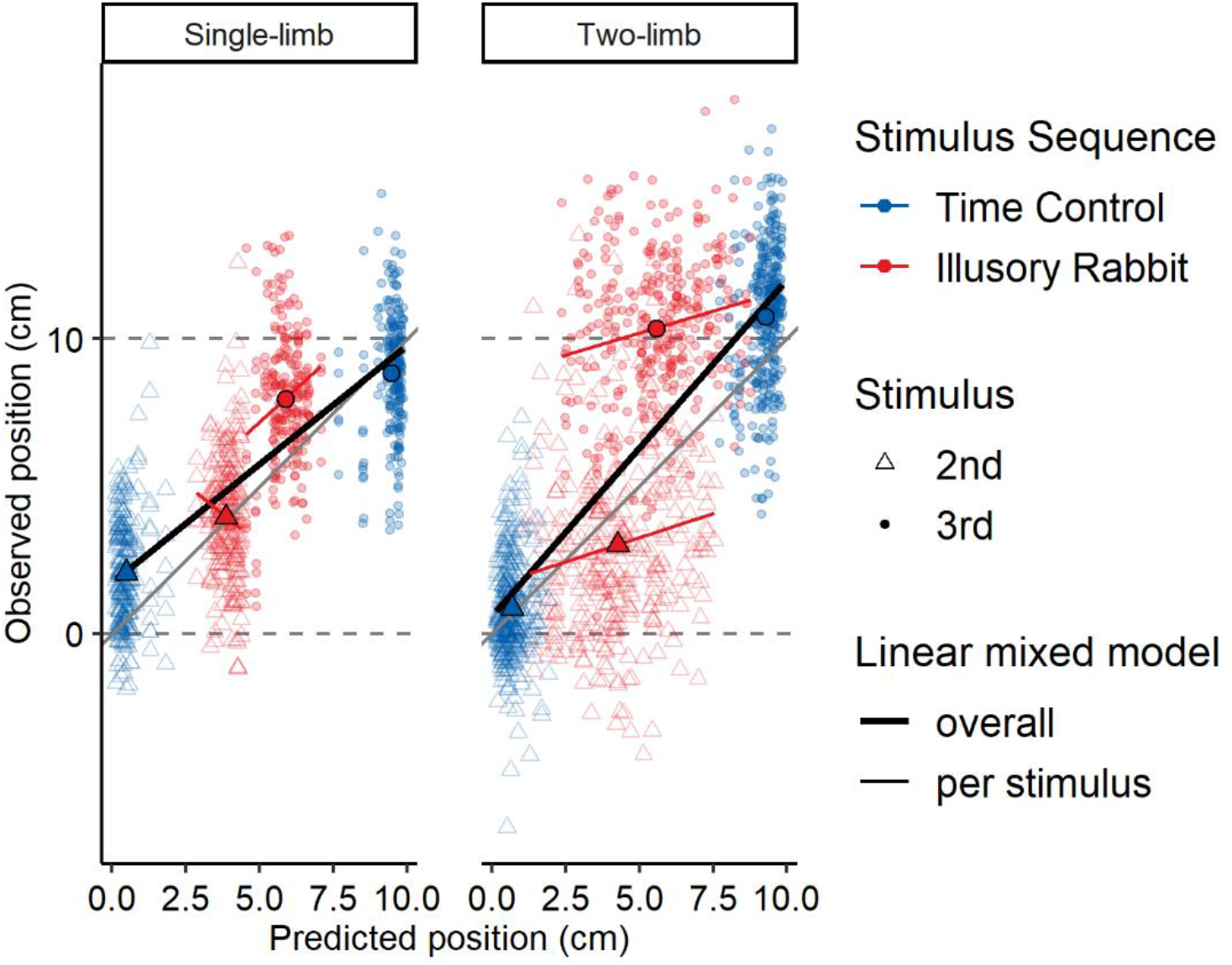
Correlations between positions that were predicted by the Bayesian observer model and observed positions. The semi-transparent points represent the mean values (averaged over participants) for stimulus patterns separately for the single-limb (Arm and Leg) and the two-limb configurations (Two Arms, Arm Leg - one side, Arm Leg - two sides). For the single-limb, all stimuli were presented within the same limb; in two-limb configurations, all patterns crossed between limbs, for the arm-leg configurations either in arm-to-leg or in leg-to-arm direction. Note that this means that each participant is represented several times in the plot as semi-transparent points; these repeated measurements are accounted for by the linear mixed models used for statistical assessment. The results of the models are shown as the regression lines. The thick black line indicates the regression over the second and third stimuli and both stimulus sequences together; the thin red lines for the illusory rabbit stimulus sequence for the second and the third stimuli separately. The thin grey diagonal line indicates perfect correspondence between model predictions and observed values. Values below this line indicate that positions are over-, values above the diagonal that they are underestimated by the Bayesian model. The larger filled symbols indicate mean values for the stimuli, allowing to see how far the averages diverge from the diagonal line.

Overall, the model made accurate predictions for both single-limb and two-limb configurations, evident from the near 1:1 relationship of predicted with observed positions in both situations. However, there were systematic discrepancies with respect to a displacement of the third stimulus towards the second stimulus, with the Bayesian model overpredicting displacement. This discrepancy was stronger in the two-limb than in the single-limb configurations.

The previous analysis assessed overall model adequacy across the two stimuli that were affected by our experimental manipulations. Next, we assessed whether differences in sensory precision across participants were consistent with the Bayesian model’s prediction separately for the second and the third stimulus. According to the model, the second stimulus should be displaced more towards the third and the third more towards the second the less precise a given participant was able to localize tactile stimuli. If this relationship held for two-limb configurations, this would provide evidence that the brain simultaneously considers individual variability (inverse precision) estimates of the different body parts. The LMM indicated that the relationship between predicted and observed locations varied for *Stimulus* (second vs. third) and *Limbs* (single-limb vs. two-limb configurations; χ²(1) = 8.51; p = .004). The slopes can be compared in Figure 11; they were significantly positive in the single-limb configurations for the third stimulus (b = .90, t(1092) = 2.86, p = .006) but not for the second stimulus (b = −.82, t(1094) = −1.77, p = .077), and in the two-limb configurations both for the second (b = .32, t(1077) = 3.68, p = .001) and the third stimulus (b = .27, t(1077) = 3.40, p = .001). The absence of a significant relationship for the second stimulus in the single-limb configurations may be related to the restricted variance in localization variability in the single-limb configurations (see Figure 11, left panel). Overall, there were mostly positive relationships between individuals’ tactile variability and their proneness to the rabbit illusion for each separate stimulus. Yet, regression slopes were clearly below 1, indicating that only a proportion of the variance in the rabbit effect can be accounted for by individuals’ sensory precision. In summary, the magnitude of the rabbit effect is related to participants’ perceptual precision even when stimulus sequences include two limbs. This result is in line with the notion of a statistical integration process based on external, Euclidian space.

## Discussion

Humans make large errors when they judge where a touch occurred on their body. In the context of tactile movement, these errors probably stem from reliance on past experience with objects moving across the body surface (Goldreich, 2007; Goldreich & Tong, 2013). Whereas previous accounts of tactile localization bias have focused on early interactions during tactile processing, for instance in the body periphery and in primary somatosensory cortex, we tested here whether tactile localization bias reflects movement trajectories in external space – consistent with the experience available when the body interacts with the world: objects in external space often move along trajectories that touch multiple body parts and do not adhere to the internal organization of the skin.

We addressed this question using the cutaneous rabbit illusion, a reliable tactile mislocalization paradigm based on apparent motion. Our study revealed three key findings. First, the cutaneous rabbit effect – a bias to report the second of three tactile stimuli as displaced towards the last stimulus of the sequence – occurred for sequences across two limbs, the two arms or an arm and a leg. In both cases, the stimulated skin regions were anatomically distant. Moreover, the effect was similar when stimuli were presented within and across the body midline, such as when we stimulated an arm and the same-side or opposite-side leg. With respect to distance on the skin, arm and leg locations are almost at a maximal possible distance for the human body. Similarly, arm and leg regions in S1 are distant as well, all the more with limbs across the two body halves. Therefore, the cutaneous rabbit effect did not critically depend on anatomical proximity of the tactile sequence. Second, the illusory location of the rabbit stimulus – the second stimulus in the sequence of three – was not bound to the limb on which it occurred. Participants frequently reported this stimulus to have occurred on the other limb. Thus, the tactile percept was sometimes wrongly assigned to an anatomically distant body part, even of the other body side. This finding suggests a surprising independence of tactile perception from the original location at which the stimulus touched the skin. Third, critical aspects of participants’ localization behavior were reproduced by a Bayesian observer model that integrated all three stimuli of the tactile sequences used in our experiments and used Euclidian distance between tactile locations – independent of which limb the stimulus had been presented to – as input. Thus, mislocalization of apparent motion stimuli is well explained by recurring on external-spatial prior experience. We will discuss these points in turn.

### The cutaneous rabbit effect reflects external-spatial priors

External space is the common denominator of all body-world interactions. In other words, if tactile localization errors in the rabbit illusion reflect prior experience with objects and events in the world, mislocalization should reflect typical external-spatial trajectories, that is, straight or near-straight Euclidian trajectories between two tactile locations. Moreover, many such straight-moving objects touch multiple body parts along their way, and therefore the rabbit illusion should not be confined to single limbs. Notably, most previous accounts of the rabbit effect have recurred to local interactions between neurons and their receptive fields in the periphery, or to local neural interactions in primary somatosensory areas (Blankenburg et al., 2006; Geldard, 1975; Geldard & Sherrick, 1983). Nevertheless, an influence of external space on the rabbit effect has been implicit in several previous studies that have demonstrated rabbit effects across arms (Eimer et al., 2005), different fingers of one hand (Trojan et al., 2014; Warren et al., 2010), and on a stick held between fingers (Miyazaki et al., 2010). Here, we demonstrated that the rabbit stimulus was systematically displaced along the Euclidian trajectory between stimuli presented on different limbs and body sides. Our results imply that the localization errors associated with the rabbit illusion are strongly affected by external space and that anatomical units such as a single limb and distance on the skin or in neuronal maps of primary somatosensory cortex do not prevent illusory mislocalization.

It may seem surprising that our results directly contradict Geldard’s (1982) original claim that the cutaneous rabbit effect does not cross the body midline. One explanation for these seemingly opposing results is the use of different localization methods. Geldard and collaborators mostly used a sectioning method (Geldard, 1975) in which participants verbally reported the degree of attraction as a proportion of the distance between the anchored outer stimulus positions. In contrast, our and other more recent studies have employed more direct localization methods such as indicating the perceived location by hand pointing (Miyazaki et al., 2010; Trojan et al., 2010) or by localizing the perceived location on a drawing (Flach & Haggard, 2006) or photo (our Experiment 1).

However, a different aspect of previous and our current results may be better able to clear up the seeming contradiction. In our study, the rabbit effect was significantly stronger, with more instances of illusory displacement, when all stimuli were presented on a single limb than when sequences were distributed across limbs. Thus, although involvement of distant skin locations did not prevent the rabbit illusion, anatomical factors supported its occurrence (Trojan et al., 2014). Thus, both external and anatomical factors appear to shape localization in touch. The idea that anatomical and external space interact has been proposed as the origin of several tactile illusions and effects (e.g. Aristotle Illusion: Hayward, 2008; crossing effect: Badde & Heed, 2016; Heed & Azañón, 2014; Shore et al., 2002; Yamamoto & Kitazawa, 2001). The influence of anatomical factors is compatible with views that attribute the rabbit effect to early unimodal somatosensory processing (Flach & Haggard, 2006; Geldard, 1985; Geldard & Sherrick, 1972). The multiple spatial influences evident in the rabbit effect, then, likely reflect the organization of the different involved systems – a 2D skin sheet on which touch is first sensed, and a 3D space in which interactions of the own body and other entities take place – and the requirement of interpreting between them to successfully master sensorimotor control (Medina & Coslett, 2010; Tamè et al., 2019).

### Assignment of touch to distant limbs

In a significant proportion of trials, participants reported the rabbit stimulus on the wrong limb. To be clear, this result implies that participants perceived stimuli, say, on the leg although they had actually been touched on the arm. We have observed this surprising finding in our two experiments, on two different samples, with two distinct methods. This incorrect assignment could not be explained by uncertainty associated with errors or the context of a perceptual illusion. These findings suggest that participants truly perceived tactile stimuli at distant limbs and that this perceptual assignment of stimuli to a limb was influenced by the external-spatial layout of the limbs in our experiment. In a previous report, rabbit effects included incorrect assignment of stimuli between fingers of a same hand (Trojan et al., 2014; Warren et al., 2010). However, we are first to report such incorrect assignment between non-homologous limbs (arm and leg), between limbs of the two body sides, and between body parts that are not neighbors or homologues in S1.

A previous study, like ours, presented tactile stimuli along the two arms but concluded that participants did not make errors in assigning stimuli to their true limb (Eimer et al., 2005). However, this conclusion was based on indirect assessment: participants judged whether the second stimulus was perceived at a predefined intermediate position or not. Therefore, participants may have felt stimuli on the incorrect arm but still have disagreed that it had occurred at the position predefined by the experiment. In the present study, in contrast, we used measures that allowed direct assessment of whether wrong limb assignment did or did not occur.

Although it seems surprising that humans report stimuli presented on an arm to have occurred on a leg, previous studies have reported evidence for such confusion of limbs. These studies used manipulations of body posture, with arms and legs crossed over the midline. For instance, when participants have to indicate which of two tactile stimuli, one on a crossed hand and one on a crossed foot, occurred first, they often report the wrong limb (Schicke & Röder, 2006). Whereas in that paradigm, participants performed a forced choice between only the two options of “hand” and “foot”, a recent study reported that participants also chose an incorrect limb – and sometimes a foot for a hand and vice versa – when they had to assign a single stimulus to one of the four limbs (Badde et al., 2019). Thus, our present results of arm/leg confusion are not without precedence and add to a growing body of evidence that the perception of where on the body a touch occurred can be categorically wrong.

Such errors in assigning touch to far distant limbs may seem dysfunctional. However, it is important to emphasize that the rabbit illusion is elicited by very unusual stimulus sequences that are designed to tickle out the dependence of perceptual processing on prior experience. In fact, there are other situations in which assigning tactile input to locations other than those at which they truly arise is highly functional: it is now well-established that humans are able to precisely localize where a tool they hold was touched (Heed, 2019; Miller et al., 2018, 2019), although, of course, the tactile input is sensed with the hand rather than the tool itself. Ultimately, the perception of touch on tools may well be forged by experience in the same way as our tactile perception on the body emerges from our lifelong interaction of objects with our skin (Miller et al., 2019). That the cutaneous rabbit is perceptually localized off the body when participants hold a pen between their stimulated fingers (Miyazaki et al., 2010) is, in our view, good evidence for such equivalence.

The existence of a low-velocity prior has been proposed in visual and auditory perception as well (Senna et al., 2015; Stocker & Simoncelli, 2006), modalities for which saltation, the general term for the rabbit effect, has been observed (e.g. Geldard, 1976; Getzmann, 2009; Khuu et al., 2010, 2011; Shore et al., 1998). The combination of visual flashes with tactile stimuli can also affect the strength of the rabbit illusion (Asai & Kanayama, 2012). That the illusion occurs in all three modalities further supports the idea that it is mapped on world-space rather than strictly anatomical coordinates.

### A Bayesian observer model fed with external distances replicates central aspects of the empirical data

The most elaborate model of tactile localization to date is the Bayesian observer model (Goldreich, 2007; Goldreich & Tong, 2013). This model predicts the perceived locations of stimuli based on localization variability and a prior that assumes single stimuli to result from a common object that slowly moves across the skin. We demonstrated that model predictions match core aspects of behavior on the level of individual participants when stimulus locations are represented in external space and the distances between them as Euclidian distances. Generally, there were strong positive relationships between predicted and observed positions indicating correspondence between the model and the data both in the single-limb and in the two-limb configurations. Therefore, the mechanisms that mediate the statistical integration of perceptual precision and a low-velocity prior are not restricted to single limbs, continuous skin stretches, or even the same cortical hemisphere. These results suggest the principal suitability of the model to prior experience made in external, Euclidian space.

Our empirical data mirrored the displacement of the second towards the third stimuli as one of the key predictions of the Bayesian observer model as elaborated above. We also observed displacement of the third towards the position of the first/second stimuli consistent both with the Bayesian model and previous experimental results (Miyazaki et al., 2010; Trojan et al., 2010) although this effect was weaker than predicted by the model .

Moreover, the predicted dependence of the rabbit illusion on spatial acuity of the touched body part (Geldard & Sherrick, 1983; Goldreich, 2007; Goldreich & Tong, 2013; Tong et al., 2016; Warren & Tillery, 2011), receives at least partial support by our modelling. We observed stronger displacements of the second stimulus in a leg-to-arm than in an arm-to-leg direction, that is, higher displacement when the stimulus was presented on a surface with lower acuity. However, this effect was only present for the arm and the leg of the same body side, suggesting that this kind of integration depends on body half-specific processing or processes within a cortical hemisphere. Further, we did not find a significant difference when the single arm and the single leg configurations were compared with each other. The reason for this latter finding is not clear. It is of note that the differences in localization variability between arm and leg were also smaller when we compared the single limb configurations as opposed to the limbs in the two-limb configurations. Possibly, participants compensated the perceptual imprecision more efficiently in single than two-limb limb configurations, potentially through attentional or expectation mechanisms, which may have diminished differences in rabbit patterns between the limbs.

Although the model reproduced key aspects of the observed patterns, there were systematic discrepancies between model predictions and observed data. First and foremost, the model predicted a stronger displacement of the third towards the second stimuli than observed in the data. A possible explanation is that the displacement of the third stimulus might be difficult to detect using the three-stimulus rabbit sequences. One study (Kilgard & Merzenich, 1995) used a four-stimulus sequence with an additional reference stimulus (at position 2) with a relatively long SOA between the third and this fourth stimulus. There, a symmetrical attraction of the second and third stimuli towards one another was evident. Presumably, the presentation of an additional reference stimulus facilitated perceiving the displacement of third towards the second stimulus. Whether the displacement of the third stimulus is also weaker than predicted by the model in previous studies that used a three-stimulus sequence (Miyazaki et al., 2010; Trojan et al., 2010) is unknown because these studies did not provide formal analyses to compare predictions to data. Based on the present study, it can be concluded that the overestimated displacement of the third stimulus in the three-stimulus illusory rabbit sequences is related to anatomical factors because the model-data discrepancy was especially strong for the two-limb configurations. Perceived displacements of the third stimulus, therefore, depended on stimulus sequences being presented along a continuous stretch of skin and, hence, indicate an anatomically based effect.

Second, we observed a stronger displacement of the second stimulus in the time control sequences than predicted by the model. It is possible that this effect reflects participants’ expectations about stimulus positions based on the statistics of trials presented in the experiment. Participants perceived location control stimuli with veridical displacements of the second stimulus and they might have built up expectations of displacements also for the time control sequences. We conclude that the Bayesian model predicted key features of the localization behavior when using external Euclidian space, but that there are patterns in the data that are not captured by the model. These patterns require additional assumptions about anatomical factors or expectations about locations beyond a simple low-velocity assumption. The fact that some model predictions were not met reverberates the above-mentioned suggestion that external-spatial priors alone do not fully capture the localization errors in the rabbit illusion paradigm. Future iterations of the Bayesian observer model could implement both external and anatomical factors to test whether localization can be better explained by integration across multiple spatial codes. At this stage, however, it is not clear how this anatomical prior could be defined. A straight-forward approach may be to integrate the Euclidian distances derived from external-spatial stimulus locations, as used in the present study, with 2D distances in skin space. Yet, the observation that the displacement of the second stimulus was comparable between the AL-1s and AL-2s configurations and the fact that it was quite accurately predicted by the model makes it unlikely that the missing anatomical prior relates to distance in 2D skin space. If the model lacked a 2D skin prior, rabbit effects should have been weaker in the AL-2s because the anatomical distance is larger in that limb configuration than in AL-1s. It is possible that the brain uses another, yet unknown prior, for example based on limb assignment (Badde et al., 2019; Maij et al., 2020), that increases the perceived distance between stimuli once the brain has inferred they had occurred on different limbs. This might be linked to recent Bayesian causal inference accounts (Debats et al., 2017; Kayser & Shams, 2015; Körding et al., 2007) which posit that the process of (multi)sensory integration has two stages: in one process the brain infers whether stimuli should be integrated or segregated; in another process the brain computes the perceived features, for example perceived positions, for the case of integration and segregation. Although this framework is usually applied to multisensory integration, it is possible that the same principles apply for the integration of multiple sources of spatial information in unimodal tactile perception. This framework could potentially also explain our finding that the illusion did not occur in all trials as resulting from an inference process of perceptual segregation.

### Limitations

#### Mislocalization in control sequences

High proportions of illusion-like responses in control sequences for the cutaneous rabbit effect have been observed before (e.g. Asai & Kanayama, 2012; Trojan et al., 2010; Warren et al., 2010). Our results showed that this phenomenon is accentuated on the leg compared to the arm. It is possible that our SOA of 700 ms was not sufficiently long to abandon the rabbit illusion completely. Note that in the study by Trojan and collaborators (2010) even at very long SOAs of up to one second there were still systematic displacements, although of smaller magnitude than observed here. Additionally, participants build up expectations about stimulus position during the course of an experiment (Brooks et al., 2019; see also Chalk et al., 2010). Whereas we view it as a strength of the experimental design that the different control conditions discourage habitual reporting rabbit-like displacements, it is, on the other hand, thinkable that illusory rabbit and location control sequences, through their middle stimulus, together biased the localization in time control sequences towards displacement.

#### Attention

Whereas we interpret the stronger rabbit effects on single limbs as reflecting the influence of anatomical factors, a hypothetical alternative is that participants could focus attention better on the stimulus array in one than two arm configurations. This is because attentional focus is known to affect the perceived position of the second stimulus (Flach & Haggard, 2006; Goldreich, 2007; Kilgard & Merzenich, 1995). Generally, we attempted to minimize the influence of cognitive factors, such as attention and expectations about stimulus locations in our design. First, the stimulated limbs were placed under a cover during the whole session and, during preparation of the tactile stimulus array; we added dummy stimulators to prevent any visual anchor (Trojan et al., 2010). Second, we randomized the direction of stimulation (from limb 1 to limb 2 and vice versa) and employed multiple sequences so that focusing on one particular position of the array would not have had a consistent effect across trials (Goldreich & Tong, 2013). Moreover, we interspersed two-limb sequences with single-limb sequences to obscure whether or not stimulation would involve several limbs. Overall, therefore, we do not consider attention a relevant factor for the result patterns we have presented.

## Conclusion

In sum, we demonstrated cutaneous rabbit effects when stimulus sequences comprised two limbs and showed that the second stimulus of three can be perceptually displaced from the truly stimulated to the other involved limb. These findings suggest that the cutaneous rabbit illusion does not primarily reflect anatomical processing in the periphery and primary somatosensory areas, but instead relies on external-spatial priors that arise from life-long tactile interactions with objects in 3D space. Key aspects of human tactile localization behavior in the rabbit illusion are accounted for by a Bayesian observer model that is fed with stimulus distance in space, rather than along the skin. Nevertheless, stronger effects for stimulus contexts that involve only a single limb suggest that anatomical factors, too, play a role in the emergence of the rabbit effect. The existence of anatomical factors is consistent with early accounts of the phenomenon, and these factors may be critical to further improve a Bayesian integration account of tactile localization.

## Acknowledgements

This work was supported by the German Research Foundation (DFG) through the Cluster of Excellence Cognitive Interaction Technology ‘CITEC’ at Bielefeld University, and an Emmy Noether grant to TH (He 6368/1-3). We thank Nele Rödenbeck and Sarah Bremsteller for their support with data acquisition, and Odile Sauzet for statistics consulting.

## Data availability statement

All data and code and digital material used in this study are available on the website of the Open Science Framework (OSF) and can be accessed at https://osf.io/a8bgt/.

## CRediT author statement

**Marie Martel:** Conceptualization, Methodology, Formal analysis, Data Curation, Writing - Original Draft, Writing - Review & Editing, Visualization. **Xaver Fuchs:** Conceptualization, Methodology, Software, Formal analysis, Data Curation, Writing - Original Draft, Writing - Review & Editing, Visualization. **Jörg Trojan:** Conceptualization, Methodology. **Valerie Gockel:** Conceptualization, Methodology, Investigation. **Boukje Habets:** Conceptualization, Methodology, Writing - Review & Editing. **Tobias Heed:** Conceptualization, Methodology, Resources, Writing - Original Draft, Writing - Review & Editing, Supervision, Project administration, Funding acquisition.

## References

Adams, W. J., Graf, E. W., & Ernst, M. O. (2004). Experience can change the “light-from-above” prior. Nature Neuroscience, 7(10), 1057–1058. https://doi.org/10.1038/nn1312

Asai, T., & Kanayama, N. (2012). “Cutaneous rabbit” hops toward a light: unimodal and cross-modal causality on the skin. Frontiers in Psychology, 3, 427. https://doi.org/10.3389/fpsyg.2012.00427

Badde, S., & Heed, T. (2016). Towards explaining spatial touch perception: Weighted integration of multiple location codes. Cognitive Neuropsychology, 33(1–2), 26–47. https://doi.org/10.1080/02643294.2016.1168791

Badde, S., Röder, B., & Heed, T. (2019). Feeling a Touch to the Hand on the Foot. Current Biology: CB, 29(9), 1491–1497.e4. https://doi.org/10.1016/j.cub.2019.02.060

Barnett-Cowan, M., Ernst, M. O., & Bülthoff, H. H. (2018). Gravity-dependent change in the ‘light-from-above’ prior. Scientific Reports, 8(1), 15131. https://doi.org/10.1038/s41598-018-33625-2

Barr, D. J., Levy, R., Scheepers, C., & Tily, H. J. (2013). Random effects structure for confirmatory hypothesis testing: Keep it maximal. Journal of Memory and Language, 68(3). https://doi.org/10.1016/j.jml.2012.11.001

Bates, D., Mächler, M., Bolker, B., & Walker, S. (2015). Fitting Linear Mixed-Effects Models Using lme4. Journal of Statistical Software. https://doi.org/10.18637/jss.v067.i01

Blankenburg, F., Ruff, C. C., Deichmann, R., Rees, G., & Driver, J. (2006). The Cutaneous Rabbit Illusion Affects Human Primary Sensory Cortex Somatotopically. PLOS Biology, 4(3), e69. https://doi.org/10.1371/journal.pbio.0040069

Brooks, J., Seizova-Cajic, T., & Taylor, J. L. (2019). Biases in tactile localization by pointing: compression for weak stimuli and centering for distributions of stimuli. Journal of Neurophysiology, 121(3), 764–772. https://doi.org/10.1152/jn.00189.2018

Brown, H., Adams, R. A., Parees, I., Edwards, M., & Friston, K. (2013). Active inference, sensory attenuation and illusions. Cognitive Processing, 14(4), 411–427. https://doi.org/10.1007/s10339-013-0571-3

Chalk, M., Seitz, A. R., & Seriès, P. (2010). Rapidly learned stimulus expectations alter perception of motion. Journal of Vision, 10(8), 2–2. https://doi.org/10.1167/10.8.2

Debats, N. B., Ernst, M. O., & Heuer, H. (2017). Perceptual attraction in tool use: evidence for a reliability-based weighting mechanism. Journal of Neurophysiology, 117(4), 1569– 1580. https://doi.org/10.1152/jn.00724.2016

Di Luca, M., & Rhodes, D. (2016). Optimal Perceived Timing: Integrating Sensory Information with Dynamically Updated Expectations. Scientific Reports, 6(1), 28563. https://doi.org/10.1038/srep28563

Eimer, M., Forster, B., & Vibell, J. (2005). Cutaneous saltation within and across arms: a new measure of the saltation illusion in somatosensation. Perception & Psychophysics, 67(3), 458–468.

Ernst, M. O., & Bülthoff, H. H. (2004). Merging the senses into a robust percept. Trends in Cognitive Sciences, 8(4), 162–169. https://doi.org/10.1016/j.tics.2004.02.002

Flach, R., & Haggard, P. (2006). The cutaneous rabbit revisited. Journal of Experimental Psychology. Human Perception and Performance, 32(3), 717–732. https://doi.org/10.1037/0096-1523.32.3.717

Fleming, S. M., & Frith, C. D. (2014). The Cognitive Neuroscience of Metacognition (Springer).

Fox, J., & Weisberg, H. S. (2011). An R Companion to Applied Regression, Second Edition. https://socialsciences.mcmaster.ca/jfox/Books/Companion/

Friston, K. (2010). The free-energy principle: a unified brain theory? Nature Reviews Neuroscience, 11(2), 127–138. https://doi.org/10.1038/nrn2787

Geldard, F. A. (1975). Sensory saltation: Metastability in the perceptual world. Lawrence Erlbaum.

Geldard, F. A. (1976). The saltatory effect in vision. Sensory Processes, 1(1), 77–86.

Geldard, F. A. (1982). Saltation in somesthesis. Psychological Bulletin, 92(1), 136–175. https://doi.org/10.1037/0033-2909.92.1.136

Geldard, F. A. (1985). The mutability of time and space on the skin. The Journal of the Acoustical Society of America, 77(1), 233–237. https://doi.org/10.1121/1.392264

Geldard, F. A., & Sherrick, C. E. (1972). The cutaneous “rabbit”: a perceptual illusion. *Science (New York*, N.Y.), 178(4057), 178–179.

Geldard, F. A., & Sherrick, C. E. (1983). The cutaneous saltatory area and its presumed neural basis. Perception & Psychophysics, 33(4), 299–304. https://doi.org/10.3758/BF03205876

Geldard, F. A., & Sherrick, C. E. (1986). Space, Time and Touch. Scientific American, 7.

Getzmann, S. (2009). Exploring auditory saltation using the “reduced-rabbit” paradigm. Journal of Experimental Psychology: Human Perception and Performance, 35(1), 289– 304. https://doi.org/10.1037/a0013026

Gibson, J. J. (1966). The senses considered as perceptual systems. Houghton Mifflin. Gibson, J. J. (1979). The ecological approach to visual perception. Houghton Mifflin.

Goldreich, D. (2007). A Bayesian perceptual model replicates the cutaneous rabbit and other tactile spatiotemporal illusions. PloS One, 2(3), e333. https://doi.org/10.1371/journal.pone.0000333

Goldreich, D., & Tong, J. (2013). Prediction, Postdiction, and Perceptual Length Contraction: A Bayesian Low-Speed Prior Captures the Cutaneous Rabbit and Related Illusions. Frontiers in Psychology, 4. https://doi.org/10.3389/fpsyg.2013.00221

Hayward, V. (2008). A brief taxonomy of tactile illusions and demonstrations that can be done in a hardware store. Brain Research Bulletin, 75(6), 742–752. https://doi.org/10.1016/j.brainresbull.2008.01.008

Hayward, V. (2015). Tactile illusions. Scholarpedia, 10(3), 8245. https://doi.org/10.4249/scholarpedia.8245

Heed, T. (2019). Tool Use: Two Mechanisms but One Experience. Current Biology, 29(24), R1301–R1303. https://doi.org/10.1016/j.cub.2019.10.062

Heed, T., & Azañón, E. (2014). Using time to investigate space: a review of tactile temporal order judgments as a window onto spatial processing in touch. Frontiers in Psychology, 5. https://doi.org/10.3389/fpsyg.2014.00076

Heed, T., Backhaus, J., & Röder, B. (2012). Integration of hand and finger location in external spatial coordinates for tactile localization. Journal of Experimental Psychology. Human Perception and Performance, 38(2), 386–401. https://doi.org/10.1037/a0024059

Kayser, C., & Shams, L. (2015). Multisensory causal inference in the brain. PLoS Biology, 13(2).

Khuu, S. K., Kidd, J. C., & Badcock, D. R. (2011). The influence of spatial orientation on the perceived path of visual saltatory motion. Journal of Vision, 11(9), 5–5. https://doi.org/10.1167/11.9.5

Khuu, S. K., Kidd, J. C., & Errington, J. A. (2010). The effect of motion adaptation on the position of elements in the visual saltation illusion. Journal of Vision, 10(12), 19–19. https://doi.org/10.1167/10.12.19

Kilgard, M. P., & Merzenich, M. M. (1995). Anticipated stimuli across skin. Nature, 373(6516), 663–663. https://doi.org/10.1038/373663a0

Körding, K. P., Beierholm, U., Ma, W. J., Quartz, S., Tenenbaum, J. B., & Shams, L. (2007). Causal Inference in Multisensory Perception. PLOS ONE, 2(9), e943. https://doi.org/10.1371/journal.pone.0000943

Körding, K. P., & Wolpert, D. M. (2004). Bayesian integration in sensorimotor learning. Nature, 427(6971), 244–247. https://doi.org/10.1038/nature02169

Longo, M. R., Azañón, E., & Haggard, P. (2010). More than skin deep: body representation beyond primary somatosensory cortex. Neuropsychologia, 48(3), 655–668. https://doi.org/10.1016/j.neuropsychologia.2009.08.022

Maij, F., Seegelke, C., Medendorp, W. P., & Heed, T. (2020). External location of touch is constructed post-hoc based on limb choice. ELife, 9. https://doi.org/10.7554/eLife.57804

Mancini, F., Bauleo, A., Cole, J., Lui, F., Porro, C. A., Haggard, P., & Iannetti, G. D. (2014). Whole-Body Mapping of Spatial Acuity for Pain and Touch. Annals of Neurology, 75(6), 917–924. https://doi.org/10.1002/ana.24179

Medina, J., & Coslett, H. B. (2010). From maps to form to space: touch and the body schema. Neuropsychologia, 48(3), 645–654. https://doi.org/10.1016/j.neuropsychologia.2009.08.017

Miller, L. E., Fabio, C., Ravenda, V., Bahmad, S., Koun, E., Salemme, R., Luauté, J., Bolognini, N., Hayward, V., & Farnè, A. (2019). Somatosensory Cortex Efficiently Processes Touch Located Beyond the Body. Current Biology: CB, 29(24), 4276–4283.e5. https://doi.org/10.1016/j.cub.2019.10.043

Miller, L. E., Montroni, L., Koun, E., Salemme, R., Hayward, V., & Farnè, A. (2018). Sensing with tools extends somatosensory processing beyond the body. Nature, 561(7722), 239–242. https://doi.org/10.1038/s41586-018-0460-0

Miyazaki, M., Hirashima, M., & Nozaki, D. (2010). The “Cutaneous Rabbit” Hopping out of the Body. Journal of Neuroscience, 30(5), 1856–1860. https://doi.org/10.1523/JNEUROSCI.3887-09.2010

Neumann, O., & Prinz, W. (Eds.). (1990). Relationships Between Perception and Action: Current Approaches. Springer-Verlag. https://doi.org/10.1007/978-3-642-75348-0

Oldfield, R. C. (1971). The assessment and analysis of handedness: The Edinburgh inventory. Neuropsychologia, 9(1), 97–113. https://doi.org/10.1016/0028-3932(71)90067-4

Peirce, J. W. (2007). PsychoPy--Psychophysics software in Python. Journal of Neuroscience Methods, 162(1–2), 8–13. https://doi.org/10.1016/j.jneumeth.2006.11.017

R Core Team. (2018). R: A language and environment for statistical computing. R Foundation for Statistical Computing, Vienna, Austria*. URL* https://www.R-project.org/.

Russell, L. (2019). emmeans: Estimated Marginal Means, aka Least-Squares Means. R package version 1.4.3.01. https://CRAN.R-project.org/package=emmeans.

Sauzet, O., Razum, O., Widera, T., & Brzoska, P. (2019). Two-Part Models and Quantile Regression for the Analysis of Survey Data With a Spike. The Example of Satisfaction With Health Care. Frontiers in Public Health, 7. https://doi.org/10.3389/fpubh.2019.00146

Schicke, T., & Röder, B. (2006). Spatial remapping of touch: confusion of perceived stimulus order across hand and foot. Proceedings of the National Academy of Sciences of the United States of America, 103(31), 11808–11813. https://doi.org/10.1073/pnas.0601486103

Senna, I., Parise, C. V., & Ernst, M. O. (2015). Hearing in slow-motion: Humans underestimate the speed of moving sounds. Scientific Reports, 5(1), 14054. https://doi.org/10.1038/srep14054

Shams, L., & Beierholm, U. R. (2010). Causal inference in perception. Trends in Cognitive Sciences, 14(9), 425–432. https://doi.org/10.1016/j.tics.2010.07.001

Shore, D. I., Hall, S. E., & Klein, R. M. (1998). Auditory saltation: A new measure for an old illusion. The Journal of the Acoustical Society of America, 103(6), 3730–3733. https://doi.org/10.1121/1.423093

Shore, D. I., Spry, E., & Spence, C. (2002). Confusing the mind by crossing the hands. Brain Research. Cognitive Brain Research, 14(1), 153–163. https://doi.org/10.1016/s0926-6410(02)00070-8

Stocker, A. A., & Simoncelli, E. P. (2006). Noise characteristics and prior expectations in human visual speed perception. Nature Neuroscience, 9(4), 578–585. https://doi.org/10.1038/nn1669

Tamè, L., Azañón, E., & Longo, M. R. (2019). A Conceptual Model of Tactile Processing across Body Features of Size, Shape, Side, and Spatial Location. Frontiers in Psychology, 10. https://doi.org/10.3389/fpsyg.2019.00291

Tong, J., Ngo, V., & Goldreich, D. (2016). Tactile length contraction as Bayesian inference. Journal of Neurophysiology, 116(2), 369–379. https://doi.org/10.1152/jn.00029.2016

Trojan, J., Heil, M., Maihöfner, C., Hölzl, R., Kleinböhl, D., Flor, H., & Benrath, J. (2014). Spatiotemporal integration of tactile patterns along and across fingers. Neuropsychologia, 53, 12–24. https://doi.org/10.1016/j.neuropsychologia.2013.10.019

Trojan, J., Speck, V., Kleinböhl, D., Benrath, J., Flor, H., & Maihöfner, C. (2019). Altered tactile localization and spatiotemporal integration in complex regional pain syndrome patients. European Journal of Pain (London, England), 23(3), 472–482. https://doi.org/10.1002/ejp.1321

Trojan, J., Stolle, A. M., Carl, A. M., Kleinböhl, D., Tan, H. Z., & Hölzl, R. (2010). Spatiotemporal integration in somatosensory perception: effects of sensory saltation on pointing at perceived positions on the body surface. Frontiers in Psychology, 1, 206. https://doi.org/10.3389/fpsyg.2010.00206

Warren, J. P., Santello, M., & Helms Tillery, S. I. (2010). Electrotactile stimuli delivered across fingertips inducing the Cutaneous Rabbit Effect. Experimental Brain Research, 206(4), 419–426. https://doi.org/10.1007/s00221-010-2422-0

Warren, J. P., & Tillery, S. I. H. (2011). Tactile perception: Do distinct subpopulations explain differences in mislocalization rates of stimuli across fingertips? Neuroscience Letters, 505(1), 1–5. https://doi.org/10.1016/j.neulet.2011.04.057

Weinstein, S. (1968). Intensive and extensive aspects of tactile sensitivity as a function of body part, sex, and laterality. In D. R. Kenshalo (Ed.), The skin senses: Proceedings of the first International Symposium on the Skin Senses, held at the Florida State University in Tallahassee, Florida (pp. 195–218). Charles C. Thomas Publishing.

Witt, J. K., & Riley, M. A. (2014). Discovering your inner Gibson: reconciling action-specific and ecological approaches to perception-action. Psychonomic Bulletin & Review, 21(6), 1353–1370. https://doi.org/10.3758/s13423-014-0623-4

Wolpert, D. M., & Ghahramani, Z. (2000). Computational principles of movement neuroscience. Nature Neuroscience, 3, 1212–1217. https://doi.org/10.1038/81497

Yamamoto, S., & Kitazawa, S. (2001). Reversal of subjective temporal order due to arm crossing. Nature Neuroscience, 4(7), 759–765. https://doi.org/10.1038/89559

