## Supplementary File for "Illusory tactile movement crosses arms and legs and is coded in external space"

May 2021 - Revised October 2021

#### Contents

**1. Methods** **2**

1.1. Table S1: Frequency of reporting a > 10% displacement of the second stimulus in the illusory rabbit trials, in the different limb configurations, when the sequence involved one or two limbs 2

---

This document is a reproducible report, created with RMarkdown. Data and code to reproduce the analyses are provided in the accompanying repository on the website of the Open Science Framework and can be accessed via the following link: https://osf.io/a8bgt/.

This document presents supplementary material that accompanies the main paper. Please see the files in the OSF repository for code and detailed results concerning the main paper.

---

### 1. Methods

#### 1.1. Table S1: Frequency of reporting a > 10% displacement of the second stimulus in the illusory rabbit trials, in the different limb configurations, when the sequence involved one or two limbs

***Table S1. Frequency of reporting a > 10% displacement of the second stimulus in illusory rabbit sequences for each limb configuration.*** *Each limb configuration is splitted depending on whether it involves only the arm, the leg or both. In one-limb configurations (A, L), all three stimuli occurred on one limb. In two-limb configurations (AA, AL-1s, AL-2s), the three stimuli were either presented distributed across the two involved limbs - which we focus on in the main paper - or all three stimuli were presented to only one of the two limbs of the respective experimental block. When the experimental block involved two limbs, but all stimuli occurred on a single limb, the sequences were identical to those of experimental blocks in which only a single limb was stimulated. Thus, this analysis ascertains that simply adding another potentially relevant limb does not significantly affect the occurrence of the rabbit illusion.Here we show that the rabbit illusion occured independent of the presence of a potentially relevant limb.*

| Nr of Stimulated Limbs | Limb Configuration | Stimulated Limb | Mean Freq | SD |
|---|---|---|---|---|
| one limb | A | Arm | 0.890 | 0.207 |
| one limb | AA | Arm | 0.824 | 0.265 |
| one limb | AL-1s | Arm | 0.852 | 0.285 |
| one limb | AL-2s | Arm | 0.810 | 0.211 |
| one limb | AL-1s | Leg | 0.830 | 0.305 |
| one limb | AL-2s | Leg | 0.785 | 0.290 |
| one limb | L | Leg | 0.865 | 0.224 |
| two limbs | AA | Arm+Arm | 0.650 | 0.322 |
| two limbs | AL-1s | Arm+Leg | 0.756 | 0.316 |
| two limbs | AL-2s | Arm+Leg | 0.633 | 0.303 |

#### 1.2. Table S2 and Figure S1: Frequency of reporting a displacement of the second stimulus when the illusion is considered present above 5% or 10% magnitude

***Table S2. Frequency of reporting displacement of the second stimulus per Limb Configuration and Stimulus Sequence, with a 10% and a 5% threshold.*** *nTrue and nFalse stand for the number of trials in which there was a rabbit percept or not, respectively.*

| Limb Configuration | Stimulus Sequence | Mean Freq - 10% | Mean Freq - 5% | SD - 10% | SD - 5% |
|---|---|---|---|---|---|
| A | Time Control | 0.483 | 0.498 | 0.354 | 0.359 |
| L | | 0.601 | 0.629 | 0.327 | 0.324 |
| AA | | 0.384 | 0.419 | 0.284 | 0.301 |
| AL-1s | | 0.467 | 0.479 | 0.372 | 0.367 |
| AL-2s | | 0.338 | 0.362 | 0.335 | 0.355 |
| A | Illusory Rabbit | 0.890 | 0.897 | 0.207 | 0.204 |
| L | | 0.865 | 0.877 | 0.224 | 0.210 |
| AA | | 0.650 | 0.676 | 0.322 | 0.323 |
| AL-1s | | 0.756 | 0.771 | 0.316 | 0.313 |
| AL-2s | | 0.633 | 0.646 | 0.303 | 0.304 |
| A | Location Control | 0.851 | 0.866 | 0.204 | 0.197 |
| L | | 0.856 | 0.859 | 0.221 | 0.221 |
| AA | | 0.801 | 0.822 | 0.170 | 0.172 |
| AL-1s | | 0.790 | 0.810 | 0.203 | 0.201 |
| AL-2s | | 0.747 | 0.762 | 0.188 | 0.193 |

| Limb Configuration | Stimulus Sequence | nTrue - 10% | nTrue - 5% | nFalse - 10% | nFalse - 5% |
|---|---|---|---|---|---|
| A | Time Control | 244 | 252 | 266 | 258 |
| L | | 280 | 293 | 195 | 182 |
| AA | | 156 | 171 | 283 | 268 |
| AL-1s | | 177 | 183 | 246 | 240 |
| AL-2s | | 118 | 127 | 274 | 265 |
| A | Illusory Rabbit | 410 | 413 | 46 | 43 |
| L | | 379 | 385 | 58 | 52 |
| AA | | 225 | 234 | 135 | 126 |
| AL-1s | | 257 | 262 | 97 | 92 |
| AL-2s | | 208 | 213 | 134 | 129 |
| A | Location Control | 274 | 279 | 48 | 43 |
| L | | 272 | 273 | 44 | 43 |
| AA | | 258 | 265 | 67 | 60 |
| AL-1s | | 239 | 245 | 66 | 60 |
| AL-2s | | 212 | 216 | 69 | 65 |

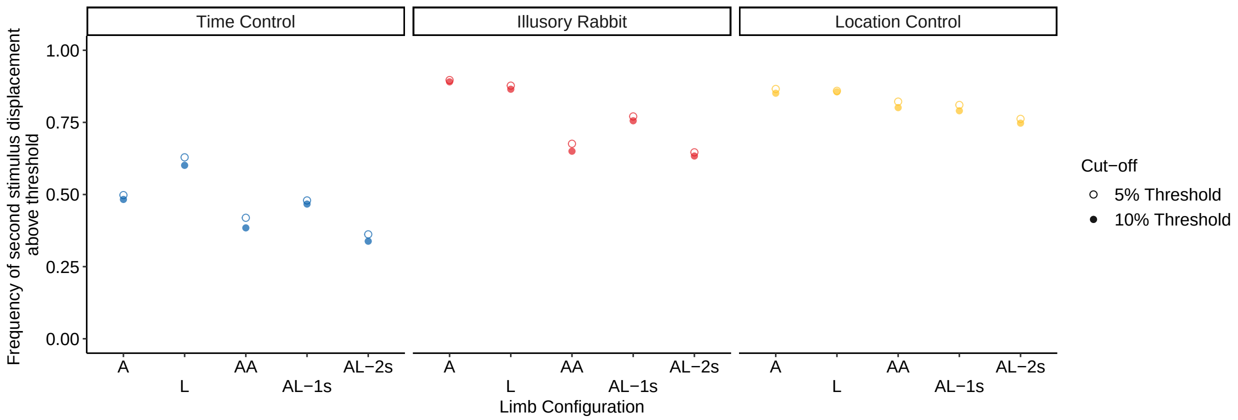

***Figure S1. Frequency of reporting a > 10% displacement of the second stimulus per Limb Configuration and Stimulus Sequence, with a 10% and a 5% threshold***

#### 1.3. Figure S2: Distribution of the magnitude of the rabbit percept for each stimulus sequence

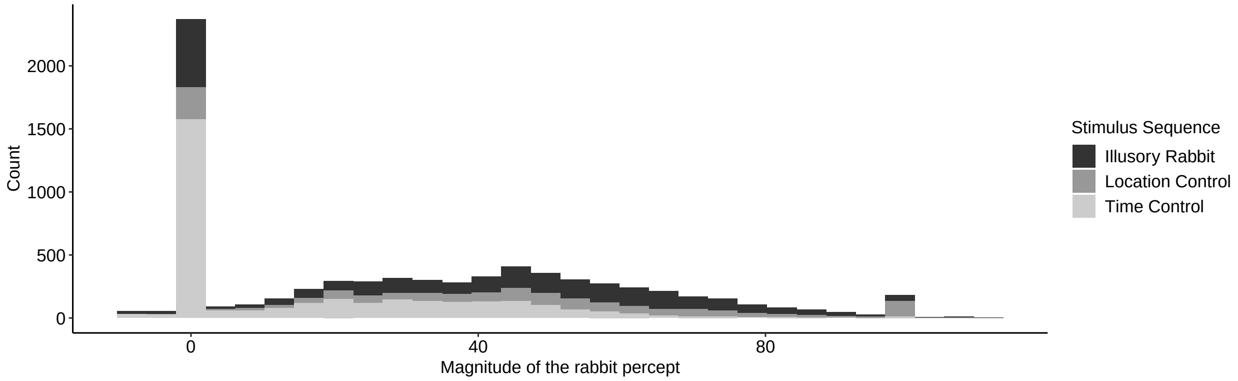

***Figure S2. Distribution of the magnitude of the rabbit percept for each stimulus sequence. Based on the distribution shown here (big proportion of values 0), we opted for two−part modelling: first we modeled the frequency of the rabbit (1/0 for presence vs. absence of the illusion), and then we remove the trials where the rabbit is absent, and modeled the magnitude of the rabbit effect.***

### 2. Experiment 1

#### 2.1. Table S3: Post-hoc comparisons of the reported positions for each stimulator in all limb configurations

***Table S3. Post-hoc comparisons of the reported positions for each stimulator in all limb configurations.*** *Values were obtained with the package emmeans (R). P values are FDR-corrected. LMM included factor Stimulators (1 to 8 or 1 to 6 depending on the limb configuration). We tested whether participants perceive distinct locations for each stimulus. To this end, we compared whether the reported localization differed significantly between each pair of stimuli. This analysis demonstrates that, indeed, localization differed for all stimulus locations.*

| limb configuration | contrast | estimate | SE | df | t ratio | p value |
|---|---|---|---|---|---|---|
| Arm (A) | 1 - 2 | -84.4 | 15.7 | 657 | -5.36 | <0.001 |
| | 1 - 3 | -199.1 | 15.8 | 657 | -12.62 | <0.001 |
| | 1 - 4 | -338.3 | 15.8 | 657 | -21.39 | <0.001 |
| | 1 - 5 | -452.4 | 15.7 | 657 | -28.73 | <0.001 |
| | 1 - 6 | -562.3 | 15.7 | 657 | -35.71 | <0.001 |
| | 2 - 3 | -114.7 | 15.8 | 657 | -7.27 | <0.001 |
| | 2 - 4 | -253.9 | 15.8 | 657 | -16.05 | <0.001 |
| | 2 - 5 | -367.9 | 15.7 | 657 | -23.37 | <0.001 |
| | 2 - 6 | -477.8 | 15.7 | 657 | -30.35 | <0.001 |
| | 3 - 4 | -139.2 | 15.9 | 657 | -8.78 | <0.001 |
| | 3 - 5 | -253.3 | 15.8 | 657 | -16.05 | <0.001 |
| | 3 - 6 | -363.1 | 15.8 | 657 | -23.01 | <0.001 |
| | 4 - 5 | -114.1 | 15.8 | 657 | -7.21 | <0.001 |
| | 4 - 6 | -224.0 | 15.8 | 657 | -14.16 | <0.001 |
| | 5 - 6 | -109.9 | 15.7 | 657 | -6.98 | <0.001 |
| Leg (L) | 1 - 2 | -68.3 | 17 | 660 | -4.01 | <0.001 |
| | 1 - 3 | -142.6 | 17 | 660 | -8.38 | <0.001 |
| | 1 - 4 | -250.0 | 17 | 660 | -14.69 | <0.001 |
| | 1 - 5 | -378.2 | 17 | 660 | -22.22 | <0.001 |
| | 1 - 6 | -443.3 | 17 | 660 | -26.05 | <0.001 |
| | 2 - 3 | -74.3 | 17 | 660 | -4.37 | <0.001 |
| | 2 - 4 | -181.7 | 17 | 660 | -10.68 | <0.001 |
| | 2 - 5 | -309.9 | 17 | 660 | -18.21 | <0.001 |
| | 2 - 6 | -375.0 | 17 | 660 | -22.04 | <0.001 |
| | 3 - 4 | -107.4 | 17 | 660 | -6.31 | <0.001 |
| | 3 - 5 | -235.6 | 17 | 660 | -13.85 | <0.001 |
| | 3 - 6 | -300.7 | 17 | 660 | -17.67 | <0.001 |
| | 4 - 5 | -128.2 | 17 | 660 | -7.53 | <0.001 |
| | 4 - 6 | -193.3 | 17 | 660 | -11.36 | <0.001 |
| | 5 - 6 | -65.1 | 17 | 660 | -3.82 | <0.001 |
| Two arms (AA): Arm 1 | 1 - 2 | -95.3 | 13 | 433 | -7.35 | <0.001 |
| | 1 - 3 | -177.4 | 13 | 433 | -13.65 | <0.001 |
| | 1 - 4 | -289.1 | 13 | 433 | -22.30 | <0.001 |
| | 2 - 3 | -82.2 | 13 | 433 | -6.32 | <0.001 |
| | 2 - 4 | -193.9 | 13 | 433 | -14.95 | <0.001 |
| | 3 - 4 | -111.7 | 13 | 433 | -8.60 | <0.001 |

***Table S3. Post-hoc comparisons of the reported positions for each stimulator in all limb configurations.*** *Values were obtained with the package emmeans (R). P values are FDR-corrected. LMM included factor Stimulators (1 to 8 or 1 to 6 depending on the limb configuration). We tested whether participants perceive distinct locations for each stimulus. To this end, we compared whether the reported localization differed significantly between each pair of stimuli. This analysis demonstrates that, indeed, localization differed for all stimulus locations. (continued)*

| **limb configuration** | **contrast** | **estimate** | **SE** | **df** | **t ratio** | **p value** |
|---|---|---|---|---|---|---|
| Two arms (AA): Arm 2 | 5 - 6 | -84.6 | 13.3 | 434 | -6.37 | <0.001 |
| | 5 - 7 | -202.1 | 13.3 | 434 | -15.20 | <0.001 |
| | 5 - 8 | -291.9 | 13.3 | 434 | -21.95 | <0.001 |
| | 6 - 7 | -117.4 | 13.3 | 434 | -8.83 | <0.001 |
| | 6 - 8 | -207.3 | 13.3 | 434 | -15.58 | <0.001 |
| | 7 - 8 | -89.8 | 13.3 | 434 | -6.75 | <0.001 |
| Arm Leg - one side (AL-1s): Arm | 1 - 2 | -83 | 10.7 | 434 | -7.75 | <0.001 |
| | 1 - 3 | -183 | 10.7 | 434 | -17.14 | <0.001 |
| | 1 - 4 | -268 | 10.7 | 434 | -24.99 | <0.001 |
| | 2 - 3 | -101 | 10.7 | 434 | -9.39 | <0.001 |
| | 2 - 4 | -185 | 10.7 | 434 | -17.24 | <0.001 |
| | 3 - 4 | -84 | 10.7 | 434 | -7.85 | <0.001 |
| Arm Leg - one side (AL-1s): Leg | 5 - 6 | -54.7 | 17.5 | 434 | -3.12 | 0.0019 |
| | 5 - 7 | -169.6 | 17.5 | 434 | -9.67 | <0.001 |
| | 5 - 8 | -230.2 | 17.5 | 434 | -13.13 | <0.001 |
| | 6 - 7 | -114.8 | 17.5 | 434 | -6.55 | <0.001 |
| | 6 - 8 | -175.5 | 17.5 | 434 | -10.01 | <0.001 |
| | 7 - 8 | -60.6 | 17.5 | 434 | -3.46 | <0.001 |
| Arm Leg - two sides (AL-2s): Arm | 1 - 2 | -65.9 | 12.5 | 434 | -5.29 | <0.001 |
| | 1 - 3 | -157.5 | 12.5 | 434 | -12.64 | <0.001 |
| | 1 - 4 | -224.7 | 12.5 | 434 | -18.03 | <0.001 |
| | 2 - 3 | -91.7 | 12.5 | 434 | -7.36 | <0.001 |
| | 2 - 4 | -158.8 | 12.5 | 434 | -12.75 | <0.001 |
| | 3 - 4 | -67.1 | 12.5 | 434 | -5.39 | <0.001 |
| Arm Leg - two sides (AL-2s): Leg | 5 - 6 | -71.0 | 15.9 | 433 | -4.46 | <0.001 |
| | 5 - 7 | -154.3 | 16.0 | 433 | -9.67 | <0.001 |
| | 5 - 8 | -257.9 | 15.9 | 433 | -16.19 | <0.001 |
| | 6 - 7 | -83.3 | 16.0 | 433 | -5.22 | <0.001 |
| | 6 - 8 | -186.9 | 15.9 | 433 | -11.73 | <0.001 |
| | 7 - 8 | -103.6 | 16.0 | 433 | -6.49 | <0.001 |

##### 2.2. Figure S3: Location of the second stimulus in trials when participants reported it on a different limb than the first (jump).

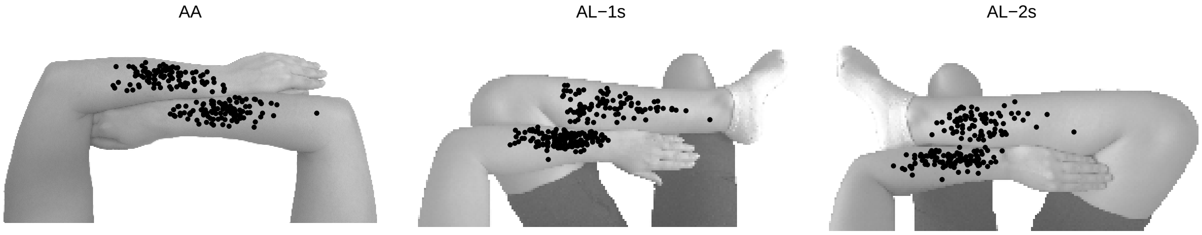

***Figure S3. Location of the second stimulus in trials when participants reported it on a different limb than the first (jump). When they reported a jump, participants localized the second stimuli well in the middle of the limb, and not at the border as a result of a mistake. They did feel the second stimuli on different limb than the first one. We included here only the two−limb conditions, when participants reported a jump.***

#### 3. Experiment 2

##### 3.1. Table S4: Post-hoc comparisons on the confidence ratings for the different stimulus sequences, when participants were correct or not

***Table S4. Post-hoc comparisons on the confidence ratings for the different stimulus sequences, when participants were correct or not.*** *Estimate indicates the differences in percent of displacement. Values were obtained with the package emmeans (R).*

| contrast | estimate | SE | df | t.ratio | p.value |
|---|---|---|---|---|---|
| Time Control,Incorrect - Illusory Rabbit,Incorrect | 1.04 | 2.17 | 79.4 | 0.479 | 0.6334 |
| Time Control,Incorrect - Time Control,Correct | -24.57 | 3.08 | 36.7 | -7.984 | <0.001 |
| Time Control,Incorrect - Illusory Rabbit,Correct | -16.44 | 3.11 | 27.8 | -5.284 | <0.001 |
| Illusory Rabbit,Incorrect - Time Control,Correct | -25.61 | 3.07 | 18.6 | -8.349 | <0.001 |
| Illusory Rabbit,Incorrect - Illusory Rabbit,Correct | -17.48 | 2.63 | 20.1 | -6.649 | <0.001 |
| Time Control,Correct - Illusory Rabbit,Correct | 8.14 | 1.40 | 20.5 | 5.795 | <0.001 |
| Illusory Rabbit,Incorrect - Location Control,Incorrect | -3.27 | 2.03 | 53.3 | -1.62 | 0.1122 |
| Illusory Rabbit,Incorrect - Illusory Rabbit,Correct | -17.27 | 2.40 | 20.3 | -7.21 | <0.001 |
| Illusory Rabbit,Incorrect - Location Control,Correct | -24.05 | 3.21 | 20.6 | -7.50 | <0.001 |
| Location Control,Incorrect - Illusory Rabbit,Correct | -14.00 | 2.49 | 24.8 | -5.62 | <0.001 |
| Location Control,Incorrect - Location Control,Correct | -20.78 | 2.60 | 27.9 | -7.98 | <0.001 |
| Illusory Rabbit,Correct - Location Control,Correct | -6.78 | 1.60 | 23.4 | -4.25 | <0.001 |

### 4. Bayesian Observer Model

#### 4.1. Table S5: Post-hoc comparisons on the precision for the two-limb configurations

***Table S5. Post-hoc comparisons on the variability for the two-limb configurations.*** *Estimate indicates the differences in variability. Values were obtained with the package emmeans (R).*

| contrast | limb configuration | estimate | SE | df | t ratio | p value |
|---|---|---|---|---|---|---|
| Limb1 - Limb2 | AA | 0.074 | 0.0777 | 54 | 0.953 | 0.3446 |
| | AL-1s | -0.852 | 0.0777 | 54 | -10.965 | <0.001 |
| | AL-2s | -0.818 | 0.0777 | 54 | -10.533 | <0.001 |

#### 4.2. Table S6: Predicted trends for each stimulus and each limb configuration

***Table S6. Predicted trends for each stimulus and each limb configuration.*** *Values were obtained with the package emmeans (R).*

| limb configuration | predicted trend | SE | df | t ratio | p value |
|---|---|---|---|---|---|
| A | 0.798 | 0.0396 | 2194 | 20.2 | <0.001 |
| L | 0.814 | 0.0425 | 2194 | 19.1 | <0.001 |
| AA | 1.209 | 0.0408 | 2194 | 29.6 | <0.001 |
| AL-1s | 1.126 | 0.0410 | 2194 | 27.5 | <0.001 |
| AL-2s | 1.090 | 0.0442 | 2194 | 24.7 | <0.001 |
